# Oxygen intrusions sustain aerobic nitrite-oxidizing bacteria in anoxic marine zones

**DOI:** 10.1101/2023.02.22.529547

**Authors:** Pearse J. Buchanan, Xin Sun, JL Weissman, Daniel McCoy, Daniele Bianchi, Emily J. Zakem

## Abstract

Anaerobic metabolisms are thought to dominate nitrogen cycling in anoxic marine zones (AMZs). However, thriving populations of aerobic nitrite-oxidizing bacteria (NOB) in AMZs challenge this assumption and remain unexplained. Using theory and modelling, we show how periodic oxygen intrusions sustain aerobic NOB in AMZs alongside more competitive aerobic heterotrophs. Ecological theory, supported by numerical simulations and genomics, frames NOB as opportunists exploiting a fleeting supply of oxygen. Consistent with *in situ* observations, simulated NOB contribute substantially to total oxygen consumption at AMZ boundaries, which implies that NOB may provide a major stabilizing feedback to AMZs. Fine-scale ocean currents increase the metabolic diversity in AMZs, which could stabilize AMZ volume under climate change.

**One-Sentence Summary:** Whiffs of oxygen to the ocean’s anoxic zones increase microbial diversity and alter biogeochemical cycling.

## Main Text

Anoxic marine zones (AMZs) form in the Eastern Pacific and Northern Indian Oceans where consumption of oxygen by respiring aerobic organisms exceeds oxygen supply. Anoxia generates a niche for anaerobic metabolisms central to nitrogen cycling and ocean-climate feedbacks. Anaerobic metabolisms remove bioavailable nitrogen, which proximally limits ocean primary productivity over half the ocean (*1*), and produce nitrous oxide (N_2_O), a potent greenhouse gas and ozone depleter (*2*). As marine deoxygenation intensifies (*3*), AMZs and nitrogen loss may expand (*4*). However, our understanding of the biogeochemical consequences of this expansion remains limited, in part because recent observations have revealed significant rates of aerobic as well as anaerobic metabolisms in AMZs (*5–10*).

How AMZs respond to ocean deoxygenation is impacted by complex and poorly understood microbial ecology. It remains unclear how anaerobic denitrification (NO_3_^-^ → NO_2_^-^ → NO → N_2_O → N_2_) and anammox (NH_4+_ + NO_2_^-^ → N_2_), which remove bioavailable nitrogen, co-occur alongside the aerobic metabolisms of ammonia oxidation (NH_3_ + O_2_ → NO_2_^-^) and nitrite oxidation (NO_2_^-^ + O_2_ → NO_3_^-^), the two steps of chemoautotrophic nitrification that remove oxygen (*9–14*). Nitrite-oxidizing bacteria (NOB) thrive in open ocean AMZs, displaying remarkably high biomass and activity in apparently anoxic waters (*12, 15–20*). Ammonia-oxidizing archaea (AOA) are also active and abundant in low oxygen waters, including the low oxygen layers above anoxic layers, where their abundance can be 10-fold higher than NOB (*8, 21–24*). However, AOA are typically less active than NOB in anoxic waters (*5, 10, 13, 16, 17*). The relative success of NOB in AMZs likely alters the rates of nitrogen loss and N_2_O production (*8, 10, 25, 26*), although the mechanism explaining their success is not yet clear. More fundamentally, thriving aerobic NOB in AMZs challenges our understanding of how diverse aerobic and anaerobic metabolisms coexist.

Multiple explanations for nitrification in AMZs have been proposed (*27*). The high oxygen affinities of both AOA and NOB suggest that these microbes can survive at nanomolar oxygen concentrations (*8, 25*), although these chemoautotrophs are unlikely to outcompete facultatively aerobic heterotrophs for oxygen (*28*). Alternatively, certain clades of AOA and NOB may have evolved nitrite oxidation pathways using electron acceptors other than oxygen (*11, 12, 27, 29*). Others suggest that episodic oxygen supply to anoxic waters might be sufficient to sustain viable populations of nitrifiers (*8, 9, 14, 30*). While these hypotheses are not mutually exclusive, a sufficient explanation must also illuminate why AOA exhibit lower activity than NOB in anoxic waters (*5, 10, 13, 16, 17*). Here, using ecological theory and modeling, we show that fleeting oxygen intrusions can explain the observations of active, thriving aerobic NOB in AMZs.

## Results

### Heterotrophs exclude aerobic nitrifiers in steady-state anoxia

A leading explanation for aerobic NOB within AMZs is their high affinity for oxygen (*8, 25*). Previous theory has shown that aerobic and anaerobic metabolisms can coexist under steady-state anoxia with a cryptic oxygen supply, consistent with observations (*6, 7, 28*). However, empirical estimates show that chemoautotrophic nitrification demands significantly more oxygen than aerobic heterotrophy (roughly 3-4 times) per mole biomass synthesized (Tables S1 and S2) (*31, 32*). Consequently, aerobic nitrifiers are thought to be outcompeted by aerobic heterotrophs when both metabolisms are limited by oxygen (*28*) Nitrifiers could be alleviated from this competitive exclusion if heterotrophs were limited by organic substrates. Affinity, oxygen demand, other traits, and ecological factors all play a role in the competition for oxygen, together setting the subsistence concentrations of the microbial populations which determine their competitive abilities (*32*).

We expanded a previous analysis (*28*) to assess the competitive outcomes of populations of the major microbial functional types within a virtual chemostat model representative of AMZ conditions (Fig. 1a; Materials and Methods). We incorporated a wider diversity of traits, accounting for differences in cell size, carbon quota, stoichiometry, maximum growth rate, biomass yields and nutrient affinities (Tables S1 and S2). We solved for the steady-state solutions across different ratios of oxygen and organic matter supply to generate an anoxic-oxic landscape (Fig. 1b).

**Fig. 1.**
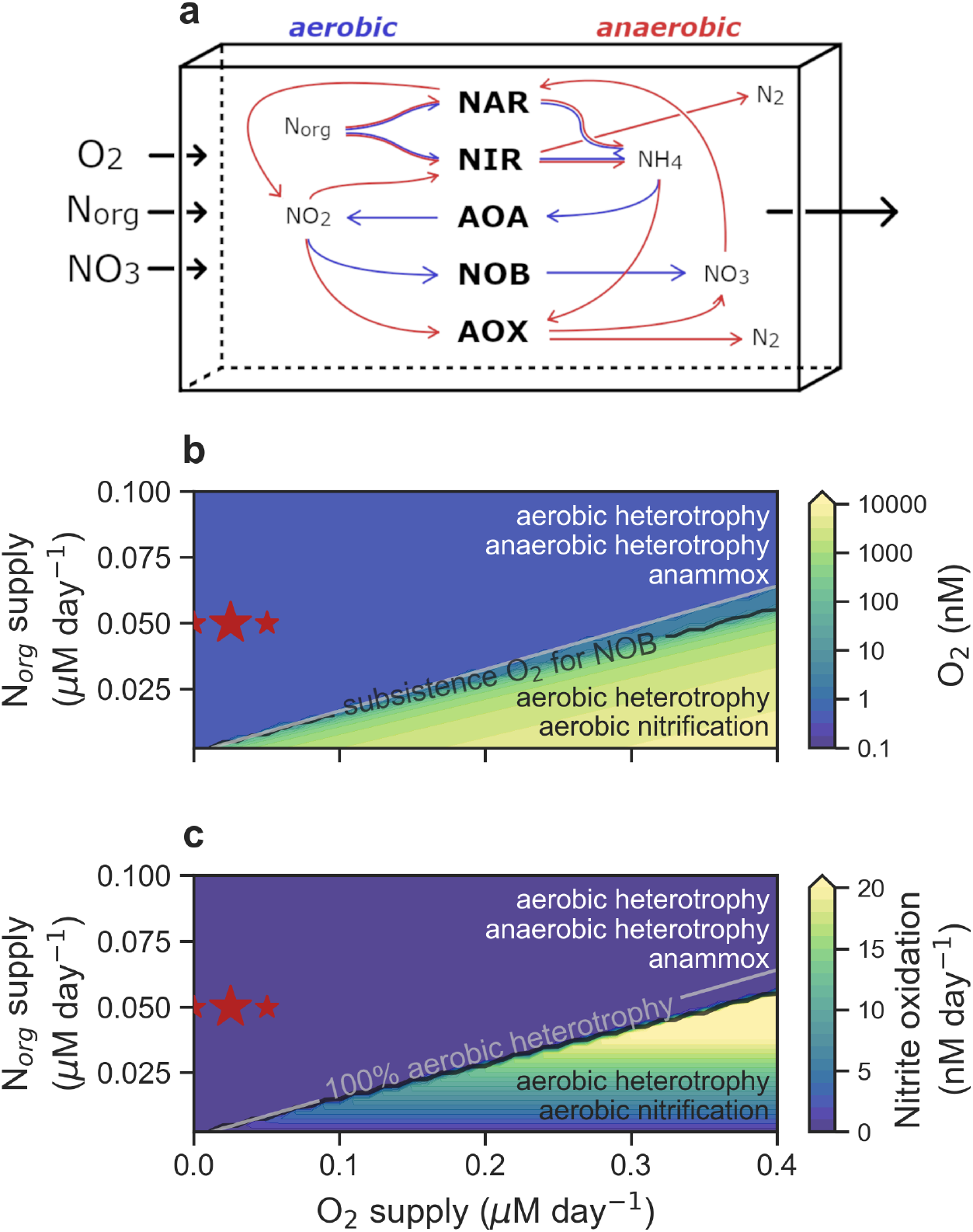
Steady-state outcomes of the virtual chemostat model. (**a**) Schematic of the model. The chemostat (single box model) contains aerobic and anaerobic metabolisms carried about by key microbes: facultative nitrate- and nitrite-reducing heterotrophs (NAR and NIR), ammonia-oxidising archaea (AOA), nitrite-oxidising bacteria (NOB) and anammox bacteria (AOX). Traits and yields are detailed in Tables S1 and S2. (**b**) Steady-state oxygen concentration as a function of different rates of organic nitrogen (N_org_) and O_2_ supply. (**c**) Rate of nitrite oxidation (nM day^-^ _1_). Silver contour marks where 100 % of heterotrophy is aerobic (following Zakem et al. (*28*)). Black contour is the subsistence O_2_ concentration for NOB (Table S3). Resulting active metabolisms in the anoxic and oxygenated regimes are listed. Red stars indicate the background (constant) N_org_ and O_2_ supply rates used in the O_2_ pulse experiments (large star shows the background rates for experiments presented in Figure 2).

Despite the diversity of traits considered, both nitrifying functional types (AOA and NOB) are consistently competitively excluded by facultatively aerobic heterotrophs when oxygen is limiting (Fig. 1c). The high affinity of AOA and NOB for oxygen (*8, 9, 25*) does enable growth at nanomolar oxygen concentrations (*10, 17, 18*) (Table S3), but heterotrophs in general are the superior competitors across the plausible trait-space because they have a lower oxygen demand for biomass synthesis (Supplementary Text; Fig. S1; Table S2). The point at which heterotrophy transitions to perform anaerobic metabolism (i.e., denitrification) occurs at oxygen concentrations that are less than the subsistence oxygen concentration required by AOA and NOB (Table S3). When considering the enormous diversity of heterotrophic bacteria, we expect that nitrifiers may outcompete a subset of inefficient aerobic heterotrophs for oxygen, while other more efficient heterotrophs will consistently exclude nitrifiers when oxygen is limiting, at least under steady-state conditions.

### Aerobic nitrifiers viable with oxygen intrusions to AMZs

The steady state can be considered as the climatological state of AMZs, set by the large-scale balance of physical supply and biological demand of oxygen. However, variability from fine-scale ocean currents sets the “weather” of the AMZs (*33*). These currents lead to time-varying intrusions of oxygen (*34*). Phytoplankton in shallow AMZ waters may also supply oxygen locally (*6*). This oxygen supply results in “cryptic” oxygen cycling, inferred from nanomolar oxygen concentrations (*35–37*) alongside non-zero oxygen consumption rates (*7, 9*). We next investigate how time-varying oxygen supply may allow NOB to thrive.

We model the biomass (*B*) of a microbial population as

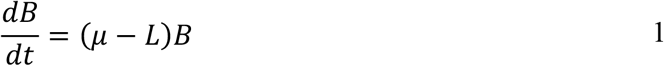

where the growth rate (*μ*) must balance the loss rate (*L*) for the population to be viable. In a time-varying environment, this balance can be achieved over time. Assuming that nitrifiers (AOA or NOB) are obligately aerobic, and neglecting limitation by micronutrients like iron (*38*), we consider oxygen intrusions to cause nitrifiers to switch from oxygen-limited growth 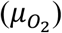 to nitrogen-limited growth (*μ*_*N*_) (Fig. 2a).

**Fig. 2.**
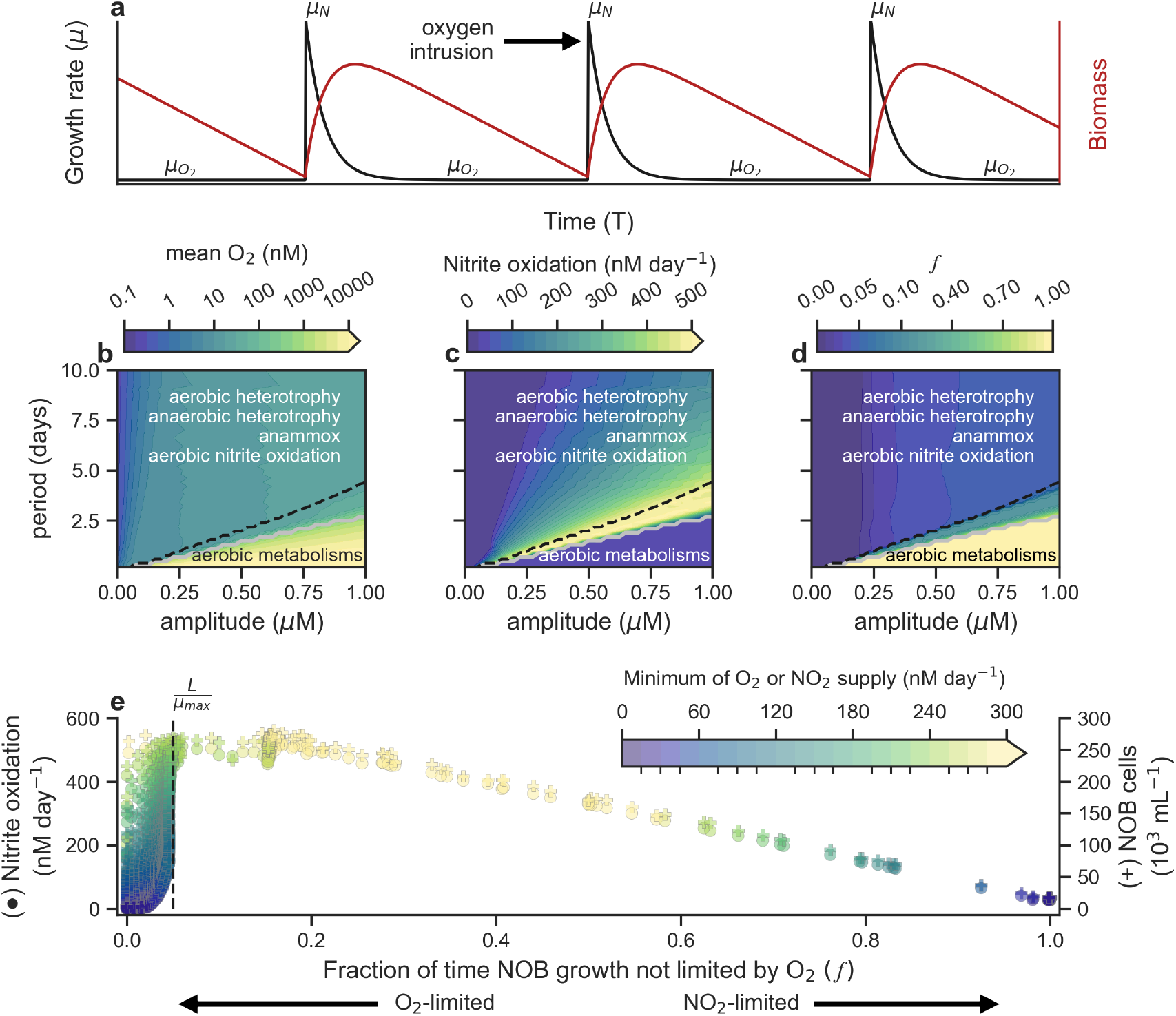
Episodic oxygen pulses support aerobic nitrite oxidation in anoxic conditions. (**a**) Schematic of oxygen intrusions to an AMZ causing a shift from an oxygen-limited growth rate 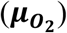 to a nitrogen-limited growth rate (***μ***_***N***_) and rapid NOB biomass accumulation (red). (**b**) Average concentration of dissolved oxygen (O_2_; nM) over last 100 days of chemostat experiments (run to dynamic equilibrium). O_2_ pulses applied over a range of amplitudes and periods (inverse of frequency). Silver contour marks where 100 % of heterotrophy is aerobic (Φ = 1, from Zakem et al. (*28*)). Active metabolisms in the anoxic and oxygenated regimes are listed. (**c**), Average rate of nitrite oxidation in nM day^-1^. (**d**) Fraction of time NOB released from oxygen limitation (***f***) in model. Black dashed line is analytically derived ***f***, where 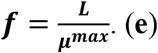 Relationship between ***f***, the rate of nitrite oxidation (circles) and NOB abundance (pluses).(*54*) Points colored by the average rate of oxygen or nitrite supply, whichever is lowest (nM day^-1^).

Following Litchman & Klausmier (*39*), we estimate the fraction of time, *f*, that nitrifiers must be released from oxygen-limited growth to sustain themselves in a low-oxygen environment (see derivation in Materials and Methods) as

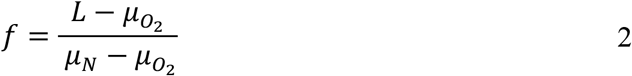

where 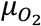 and *μ*_*N*_ are the oxygen- and nitrogen-limited growth rates, respectively, and *L* represents the integrated loss to predators, viral lysis, cell maintenance, and other mortality.

To simplify, we can assume 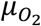 is negligible when oxygen is inaccessible (ignoring potential anaerobic metabolism (*11, 12, 27*)). Also, if the nitrogen resource (nitrite for NOB or ammonium for AOA) is abundant, then *μ*_*N*_ may approach the maximum growth rate (*μ*^*max*^). With both assumptions,

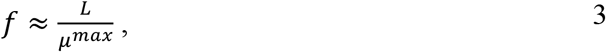

and the fraction of time that oxygen does not limit growth is approximated by the ratio of the population’s loss rate to maximum growth rate. If *L* ≪ *μ*^*max*^, *f* approaches zero, implying viability in environments with infrequent oxygen intrusions.

This theory can explain the divergent activities of NOB and AOA in AMZs. It is typical for nitrite to accumulate and for ammonium to be depleted in AMZs (*12, 30*). This difference in electron donor availability means that during an oxygen intrusion AOA cannot maximize growth (*μ*_*N*_ < *μ*^*max*^) and require more frequent oxygen intrusions to sustain a viable population (Fig. S2). Meanwhile, NOB may grow rapidly on the abundant nitrite pool.

### Chemostat simulations with oxygen intrusions support theory

Next, we applied pulses of oxygen to the virtual chemostat model to examine whether the theory is borne out when considering the AMZ microbial ecosystem (Fig. 1a). We periodically increased the input of oxygen into the chemostat, widely varying both the intensity and the frequency of the pulses (Materials and Methods). We applied these pulses to background conditions (i.e., constant supply rates of organic matter and oxygen) that generate deep anoxia and exclude nitrifiers under steady state (red stars in Fig. 1). Following an oxygen pulse, most experiments return to functional anoxia, containing ≈10 nM of oxygen when averaged in time (Fig. 2b, S3), roughly matching the subsistence oxygen concentration required to sustain a viable population of NOB (Table S3). Even large oxygen pulses are rapidly consumed by the microbial community, returning the system to long periods of oxygen scarcity beneath 10 nM (Fig. S3).

As predicted, the episodic pulses sustain a population of obligately aerobic NOB in the model. Mean nitrite oxidation rates are substantially larger under anoxic regimes with fleeting oxygenation, reaching values as high as 500 nM day^-1^, than under fully oxic or fully anoxic regimes (Fig. 2c). Peak NOB activity and biomass co-occur with *f* values close to the analytically derived *f* of Eq. 3 (dashed black line in Fig. 2b-e), indicating that coexistence of aerobic NOB with anaerobic metabolisms is beneficial. In this case, *f* is an emergent property of the model (Fig. 2d), calculated as the time-averaged growth limitation by oxygen (0) or nitrogen (1). Resulting *f* near zero indicates persistent oxygen limitation, while *f* near one indicates fully oxygenated experiments with nitrogen limitation. This nitrogen limitation occurs because nitrite does not accumulate in the absence of heterotrophic nitrate reduction, even when ammonia oxidation is active. Thus, transient oxygenation maximizes the supply of both resources (Fig. 2e).

These results are qualitatively robust across widely varying values of maximum growth rate (*µ*^*max*^) and conditions, including when *µ*^*max*^ of NOB is lower than that of heterotrophs and AOA (Fig. S4-6). As Eq. 3 suggests, a lower *µ*^*max*^ for NOB increases the frequency or strength of the oxygen intrusions required to sustain the population, shifting *f* to higher values. Meanwhile, aerobic AOA is sustained at minimal rates due to competition with anaerobic ammonium oxidation (anammox) for a limited supply of both ammonium and oxygen (Fig. S7). Our sensitivity studies, however, indicate that under organic-mater-rich conditions, faster-growing strains of AOA could theoretically replace slower-growing strains of NOB as the dominant oxygen consumers (Fig. S6). Further experimental studies are required to determine whether this theoretical outcome is realized, perhaps nearby organic particles or in highly productive coastal areas.

### Nitrite oxidizers in a 3-D model AMZ

We next investigated whether NOB thrive in AMZs with realistic ocean dynamics, where mesoscale eddies supply oxygen variably over space and time. We incorporated our microbial ecosystem model into an eddy-resolving, three-dimensional model of the Eastern Tropical South Pacific, which contains one of the major AMZs of the world ocean. Specifically, we integrated the key heterotrophic and chemoautotrophic microbial functional types (Fig. 1a) into the Biogeochemical Elemental Cycle (BEC) model (*40*), which was coupled online to the Regional Ocean Model System (ROMS) configured at an eddy-resolving (0.1° horizontal) resolution (*41*) (Materials and Methods). The model explicitly simulates the time-varying nature of oxygen intrusions within the emergent AMZ.

The model produces an AMZ offshore of the Peruvian upwelling, reaching a maximum vertical thickness of about 500 m, with nitrite accumulating. Because the model regularly generates mesoscale eddies of 10-100 km radii (*42, 43*), oxygen intrusions are common at the boundaries of the simulated AMZ (Fig. 3a). As predicted by our theory, NOB biomass and activity peak where the supply of limiting resources, oxygen and nitrite, is maximized (Fig. 3b). NOB activity is particularly high at the upper AMZ boundary where high rates of nitrate reduction, driven by a rich organic matter flux, coincide with more frequent oxygen intrusions (Fig. 3d). We provide animations of daily mean output in the Supplementary Materials.

**Fig. 3.**
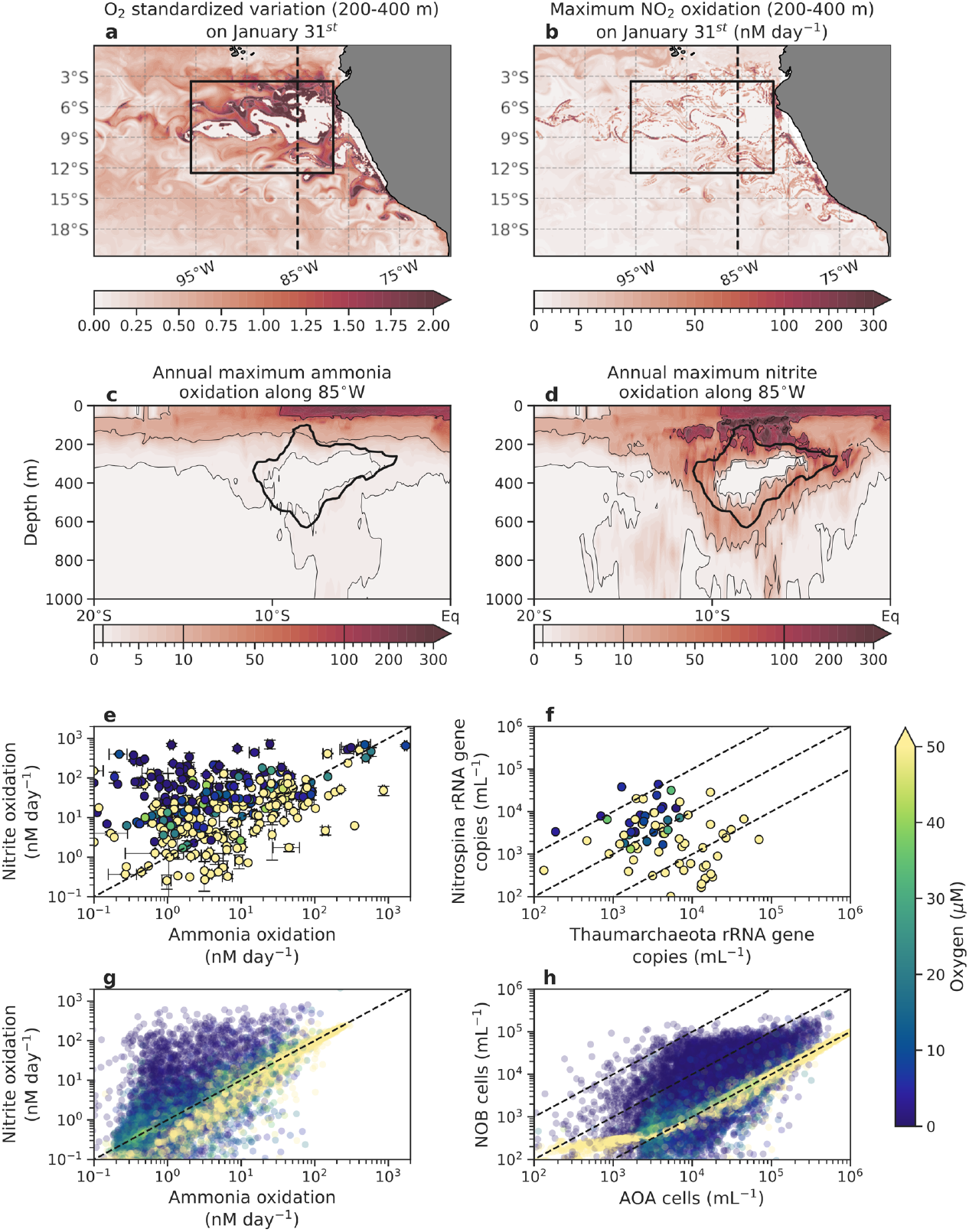
AMZ nitrification dynamics in an eddy-resolving 3D ocean model. (**a**) Standardized variation (standard deviation divided by mean) in oxygen (O_2_) between 200 and 400 meters depth on a single day. (**b**) Maximum rate of nitrite oxidation between 200 and 400 meters depth on the same day. Annual maximum values of (**c**) ammonia oxidation and (**d**) nitrite oxidation along 85ºW (dashed line in panels **a** and **b**). (**e**) Observed (*62*) and (**g**) modelled co-occuring rates of ammonia oxidation and nitrite oxidation, colored by *in situ* oxygen concentration. (**f**) Observed Thaumarchaeota (AOA) and *Nitrospina* (NOB) rRNA gene copies (*62*) and (**h**) modelled AOA and NOB cell number estimates, calculated from carbon biomass using measured cell carbon quotas of marine organisms (Table S1; Materials and Methods). Model data in panels (**g**) and (**h**) are from 2000 randomly sampled depth profiles (0-1000 m) within the box depicted in panels (**a**) and (**b**). Model statistics are generated from daily averages over the final simulation year.

While AOA is active within the simulated AMZ, AOA do not thrive to the same extent as NOB (Fig. 3c). This captures the observed decoupling between AOA and NOB in oxygenated versus oxygen-depleted environments. In oxygenated environments, ammonia and nitrite oxidation rates are of similar magnitude, and estimated AOA:NOB abundance is of order 10:1 (*44*) (Fig. 3e,f), as captured by the model (Fig. 3g,h). In contrast, in oxygen-depleted waters, nitrite oxidation rates and NOB abundances are significantly higher in both data and model.

### Aerobic NOB may stabilize anoxia

NOB contribute substantially to the oxygen consumed at the boundaries of the AMZ (Fig. 4a and S8). When integrated within the AMZ (oxygen < 1 µM), and across simulations that vary the relative *µ*^*max*^ values of the microbial functional types (Supplementary Text, Fig. S9-S11, Table S4, S5), nitrite oxidation contributes 24% to 49% of total oxygen consumption (Fig. 4b). At individual locations, this contribution varies over a wider range (Fig. 4c): the modeled interquartile range of 0% to 66% is consistent with measured contributions of 0% to 69% (Fig. 4b, Fig. S12). In the model, these location-specific contributions are low when total oxygen consumption is low (Fig. S8). Because of this positive correlation, the integrated contribution exceeds the mean of the location-specific values. Hence, an empirical estimate of NOB’s contribution to oxygen consumption across an AMZ likely requires weighting the mean of measured contributions at individual locations by the total oxygen consumption.

**Fig. 4.**
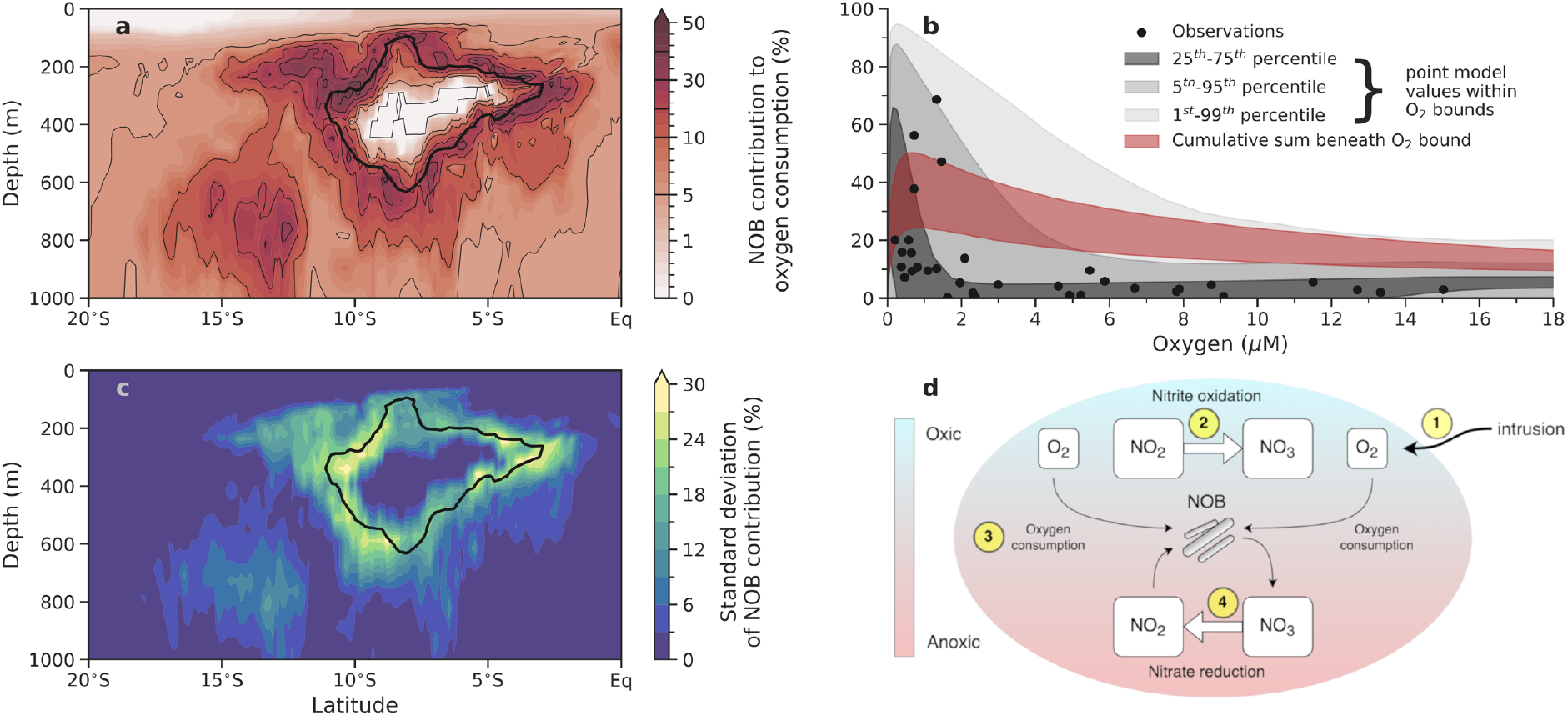
NOB contribution to oxygen (O_2_) consumption and its stabilizing feedback to oxygen perturbations. (**a**) Annual average contribution of NOB to oxygen consumption along 85ºW. The black line shows where annual mean O_2_ = 1 µM. (**b**) Observed (*9*) and modelled NOB contribution to total O_2_ consumption at different ambient O_2_ concentrations. The modelled percentiles (grey shading) refer to the NOB contribution to O_2_ consumption at individual locations, and within discrete O_2_ bounds, and thus are directly comparable with the observations. The integrated contribution (red shading) is the total NOB contribution to O_2_ consumption, cumulatively calculated beneath an O_2_ concentration. The range incorporates temporal variations and different assumptions of microbial maximum growth rates (Materials and Methods; Supplementary Text). (**c**) Temporal variability in the contribution of NOB to O_2_ consumption. All model data is generated from daily averages over the final simulation year. (**d**) Schematic of the NOB feedback to perturbations in O_2_ supply: O_2_ intrusions (1) stimulate nitrite oxidation (2) and gains in NOB biomass, which in turn elevates O_2_ consumption (3) and a more rapid return to anoxia, thereby supporting production of nitrite via anaerobic nitrate reduction (4), which maintains the ability to buffer O_2_ intrusions.

A consistent result of both the chemostat and 3-D models is that aerobic NOB show increased biomass and activity when exposed to stronger and more frequent oxygen pulses, so long as anoxic conditions return. By consuming more oxygen with more biomass, NOB exert a negative feedback to increasing oxygen supply. This sustains the anaerobic niche, which also ensures nitrite supply via heterotrophic nitrate reduction (Fig. 4d). Thus, NOB may stabilize the size and intensity of AMZs and may do so in a highly responsive manner, since their growth is not typically limited by their electron donor, nitrite, which accumulates in AMZs (*12, 30*). Heterotrophic bacteria and AOA cannot exploit an oxygen intrusion as rapidly as NOB if their growth is limited by organic matter and ammonium, respectively, which is thought to generally be the case (*45–47*), though this may differ in organic-matter-rich environments (Fig. S6). Overall, our results imply that if the oxygen supplied to AMZs declines as expected (*33*), NOB biomass and aerobic nitrite oxidation may also decline, which would in turn reduce biological oxygen demand. In this way, NOB may buffer oxygen perturbations to AMZs.

### Genome-based maximum growth rates support theory

Our results imply that the NOB inhabiting AMZs (*25, 48*) survive due to an opportunistic “boom-and-bust” life-history strategy (*39*), “booming” when oxygen is available. Furthermore, Eq. 3 suggests that NOB are better adapted to AMZs when their maximum growth rates (*μ*^*max*^) are higher. Thus, selection for faster growth to facilitate colonization deeper into AMZs, where oxygen intrusions are rarer (*49*) may leave discernable patterns of growth optimization on NOB genomes.

As a first test of this hypothesis, we employed codon usage bias statistics (*50*) to estimate *μ*^*max*^ from available metagenome-assembled genomes (MAGs) of marine NOB and other functional types (Materials and Methods, Table S6). Codon usage bias in ribosomal proteins reflects a universal signature of optimization for rapid growth (*51*). It has been shown to accurately predict growth potential even when extrapolating across phyla (*50*). However, the limited training dataset for the algorithm is biased towards heterotrophic bacteria, and so, we updated the training dataset with measured *μ*^*max*^ from 3 AOA and 6 NOB species (Fig. S13; Table S7) to improve the predictions.

The metagenome-based estimates of *μ*^*max*^ for NOB from AMZs, as well as from oxygenated environments, are higher than those of AOA and the available MAGs of heterotrophic bacteria extracted from AMZs (*Candidatus* Pelagibacter, a member of the ubiquitous SAR11 clade, known to grow slowly) (Fig. 5). Measurements from cultures (red markers in Fig. 5) also suggest higher *μ*^*max*^ for NOB than AOA. The genomic estimates lie at the high end of the measurements, in part because the estimates represent a theoretical maximum rate that may not be captured by laboratory conditions. That said, much uncertainty lies in the genomic estimates, though some confidence can be garnered from a previous clustering analysis (*50*) that identified a threshold delineating faster-growing copiotrophs, with more reliable estimates, from slower-growing oligotrophs (*39*). (This threshold depends on the assumed optimal growth temperature and is not subject to the 5-hour doubling time threshold presented in Weissman et al. (*50*)) (Materials and Methods). Many NOB *μ*^*max*^ estimates appear near or above this threshold (black dashed line in Fig. 5). Acknowledging the uncertainties, we speculate that some NOB populations may be able to grow faster than AOA and Pelagibacter in AMZs, and perhaps be considered as copiotrophs.

**Fig. 5.**
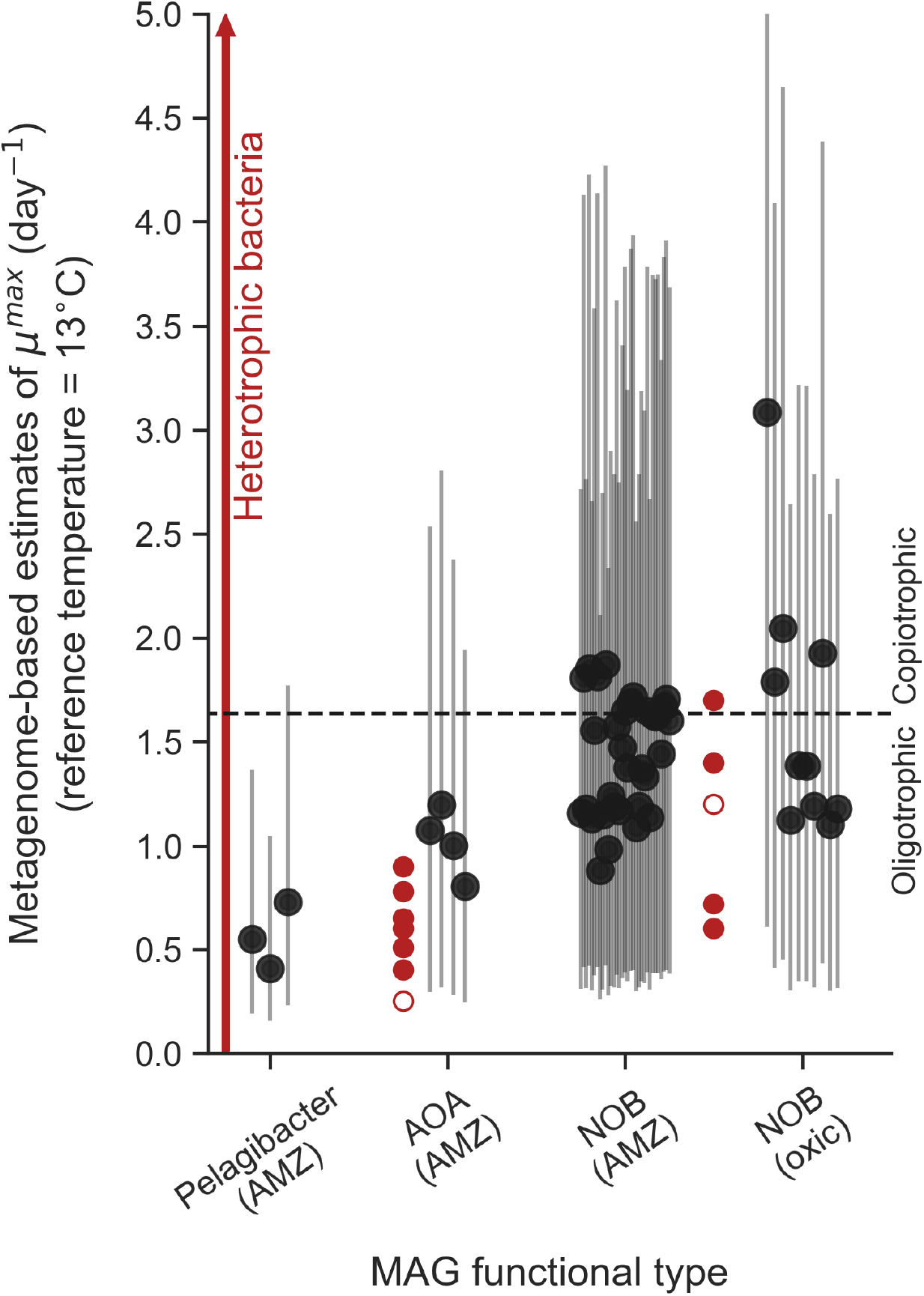
Metagenome-based estimates of maximum growth rates of marine microbial functional types. Each black dot is the central µ^max^ estimate for an individual metagenome-assembled genome (MAG) using codon usage bias statistics (*50*) for a reference temperature of 13ºC. The vertical lines are 95% confidence intervals. MAG metadata is listed in Table S6. Dashed line demarcates the threshold delineating ‘oligotrophs’ from ‘copiotrophs’ as well as the reliability of the estimates according to previous analysis (*50*). Red filled markers are measurements of µ^max^ from cultures of AOA (*38, 63–67*) and NOB (*68–70*). Red empty markers are *in situ* estimates (*24*). The vertical red arrow indicates the large range in µ^max^ for diverse heterotrophic bacteria (*71*).

This analysis supports our hypothesis that opportunistic traits would benefit NOB clades in AMZs. However, we emphasize that our main conclusion – that periodic oxygen intrusions explain the success of NOB within AMZs – is not contingent on NOB having a high *μ*^*max*^ relative to other aerobic types.

### Outlook and implications

Ocean-going studies have identified thriving populations of putatively aerobic NOB co-occurring with anaerobic metabolisms in all known major ocean AMZs (*10–14, 19, 20, 48*), leading to numerous hypotheses to explain this counter-intuitive phenomenon. Our theoretical framework suggests that AMZs prone to oxygen intrusions are in fact ideal habitats for aerobic NOB. A “goldilocks” zone of oxygen and nitrite supply exists for NOB at the oxic-anoxic boundary (*26*), a boundary that is greatly expanded by considering the folding effect of eddies (e.g., Fig. 3a). At the shallow edge of AMZs, NOB may also access diurnally pulsed oxygen supply from phytoplankton (*6*). Whether biological or physical in origin, we propose that episodic oxygen supply is key to the vigorous cycling between nitrite and nitrate observed in marine AMZs (*12, 15–19*). Though likely outcompeted by many heterotrophs in steady-state anoxia, NOB can sustain themselves by rapidly growing during oxygen intrusions, which offsets their losses during oxygen scarcity. We provide initial genomic evidence that this “boom-bust” strategy has selected for opportunistic traits in marine NOB clades.

We have here explained the elevated NOB activity and biomass in AMZs assuming an obligate aerobic metabolism. While growth via alternative electron acceptors (*11, 12, 27*) would be complementary, we argue that it is not essential where oxygen intrusions occur, even when rare. Anaerobic growth would, however, increase NOB viability, making NOB even less reliant on oxygen intrusions (Fig. S14). Additionally, NOB may also reverse the nitrite oxidoreductase enzyme and perform heterotrophic nitrate reduction (*52–54*), generating the very nitrite they require for bursts of growth during an oxygen intrusion. These anaerobic metabolisms, if accessible, would further consolidate AMZs as optimal environments and allow colonization deeper into the AMZ core.

The time-varying circulation set by mesoscale eddies is critical for understanding the presence of aerobic NOB and aerobic metabolisms in general within AMZs. For the first time, we have explicitly resolved both the mesoscale and microbial populations that drive marine nitrogen cycling within a 3-D simulated AMZ, allowing for the coexistence of aerobic NOB with anaerobic metabolisms. These *in silico* results corroborate *in situ* measurements of significant oxygen consumption by NOB (*9*), and also identify NOB as opportunistic stabilizers of anoxia, particularly at the AMZ boundaries. In this sense, NOB may in fact stimulate nitrogen loss in AMZs, a view that challenges their proposed role as suppressors (*8, 10, 25*). However, whether NOB stimulate or suppress nitrogen loss, and whether they alter the balance of anammox to denitrification and thereby affect N_2_O fluxes, will also depend on processes not well understood or resolved in this study, such as the full suite of denitrification modularity, the roles of obligately aerobic or anaerobic heterotrophic populations (*55*), and ecological interactions within particle micro-environments (*56*). These are significant challenges for our growing understanding of the nitrogen cycle. What is perhaps clearer is that the small-scale ocean circulation, the “weather” of the ocean, is essential to projections of AMZ volume and their biogeochemical implications, not only because it better represents the dynamics underpinning physical oxygen supply (*33, 57*), but also the ecological dynamics governing oxygen demand.

## Acknowledgments

We thank Clara Fuchsman, Michael Beman, Samantha Fortin and Bess Ward for sharing data and Elena Litchman, Chris Klausmier and Alessandro Tagliabue for discussions. Simulations with the 3D ocean model were run on the Expanse system at the San Diego Supercomputer Center. The authors wish to acknowledge use of the Ferret program (http://ferret.pmel.noaa.gov/Ferret/), climate data operators (https://code.mpimet.mpg.de/projects/cdo/), NetCDF Operators (http://nco.sourceforge.net/), Python (www.python.org) and GIMP (https://www.gimp.org/) for the analysis and graphics in this paper.

## Funding

Simons Foundation postdoctoral fellowship in Marine Microbial Ecology (XS)

U.S. National Science Foundation grant OCE-1847687 (DM, DB)

U.S. National Science Foundation grant 2125142 (EJZ)

Advanced Cyber infrastructure Coordination Ecosystem: Services and Support (ACCESS) program allocations TG-OCE170017 (DM, DB) and EES220053 (PJB, EZ), which are supported by National Science Foundation grants 2138259, 2138286, 2138307, 2137603, and 2138296.

## Author contributions

Conceptualization: PJB, EJZ

Methodology: PJB, EJZ, JLW, DM

Software: PJB, EJZ, DM, DB, JLW

Validation: PJB, EJZ

Formal Analysis: PJB, JLW

Investigation: PJB, XS, EJZ

Resources: EJZ, DB, JLW, XS

Data curation: PJB, XS

Writing – original draft: PJB, EJZ

Writing – reviewing and editing: PJB, EJZ, XS, DM, DB, JLW

Visualization: PJB, JLW

Funding acquisition: EJZ, DB

Project administration: EJZ

Supervision: EJZ, DB

## Competing interests

Authors declare that they have no competing interests.

## Data and materials availability

All data and code are freely available. The chemostat model, model output, data, and scripts to reproduce the figures can be downloaded at https://zenodo.org/records/15139307 (*58*). The code for the chemostat model is housed at https://github.com/pearseb/microbial_chemostat_model (*59*). The source code and the developments made to the ROMS-BEC2 ocean-biogeochemical ocean model is maintained at https://github.com/pearseb/BEC2-microbes (*60*) and analysis of ROMS-BEC2 output is held at https://github.com/pearseb/Oxygen_intrusions_NOB_ROMS_analysis (*61*).

## Supplementary Materials

Materials and Methods

Supplementary Text

Figs. S1 to S14

Tables S1 to S8

References (*62–111*)

Movies S1 to S9

## Supplementary Materials

### Materials and Methods

We developed both theoretical and numerical means of testing the effect of oxygen intrusions on the marine microbial ecological dynamics that play out in anoxic conditions. Below, we outline these approaches and discuss the supporting data.

#### Derivation of *f*, the fraction of time NOB must be released from oxygen limitation

The growth rate *μ* (day^-1^) of a given nitrifier population can be expressed as a function of the limitation to growth by either nitrogen (*μ*_*N*_) or oxygen 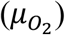, assuming Liebig’s law of the minimum, as:

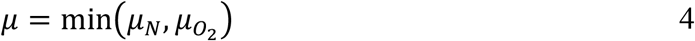

When neglecting the impacts of physical transport, the biomass (*B*; mol N m^-3^) of the population changes according to *µ* and its loss rate *L* (day^-1^), which in the environment reflects the impacts of viral lysis, grazing, and other mortality (Eq. 1). In the chemostat model, *L* is set by the dilution rate. For the population to be sustained in its environment, growth must balance losses. In a time-varying environment, this balance can be found over a sufficiently long interval of time (*I*) so that the integrated growth balances the integrated losses over time as:

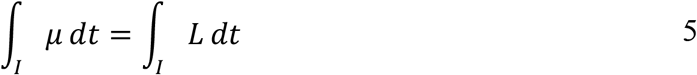

In a patchy environment with random intrusions of oxygen there may be multiple phases of growth controlled by oxygen limitation interspersed by growth controlled by nitrogen limitation (Fig. 2a). Following Litchman & Klausmier (*39*), we can decompose the integrated growth rate into two aggregate sub-intervals, one interval of N limitation *I*_*N*_ and one interval of O_2_ limitation 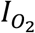, which need not be continuous in time, as:

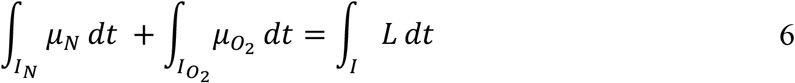

Then we define *f* as the fraction of time during interval *I* where nitrifiers are limited by nitrogen (i.e., released from oxygen limitation), and (1 − *f*) is the fraction of time where they are limited by oxygen. Neglecting any time dependence of loss rates and integrating over time gives:

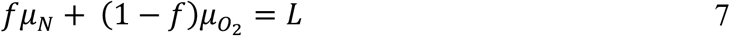

Solving for *f* gives the fraction of time that the nitrifying population can be released from limitation by oxygen to sustain itself in the environment as:

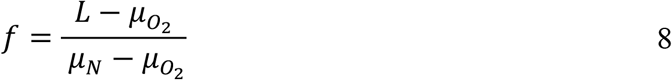

#### Zero-dimensional chemostat model

We use a zero-dimensional, theoretical model resolving multiple microbial functional type populations carrying out distinct nitrogen-cycling metabolisms (*28*). The metabolic functional types include two types of facultative heterotrophs that use oxygen for aerobic heterotrophy and either nitrate (NAR) or nitrite (NIR) as electron acceptors for anaerobic heterotrophy, anaerobic ammonium oxidation by anammox bacteria (AOX), aerobic ammonia oxidation by ammonia-oxidizing archaea (AOA), and aerobic nitrite oxidation by nitrite-oxidizing bacteria (NOB). Each metabolism requires two resources, an electron donor (S) and an electron acceptor (X), and each is performed by a microbe with unique traits (Table S1). These traits govern the competitive ability of the microbe for the resources on which they depend and the rate at which their biomass accumulates.

The basic economy of the model is in nitrogen, such that biomass is resolved in units of mol N_bio_ m^-3^. Growth is the product of the specific uptake rate of a resource (V, mol resource (mol N_bio_)^-1^ day^-1^) multiplied by the biomass yield associated with that resource (*y*, mol N_bio_ (mol resource)^-1^). The realized growth rate (*µ*, day^-1^) is the minimum of growth on S or X as:

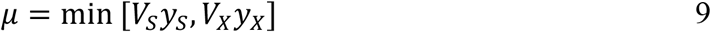

The specific uptake of a resource (*V*_*S*_, *V*_*X*_), except oxygen, is calculated via Michaelis Menten kinetics using a maximum specific uptake rate 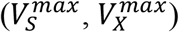 and a half-saturation coefficient (K_S_, K_X_) as:

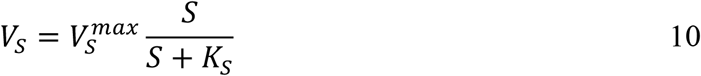

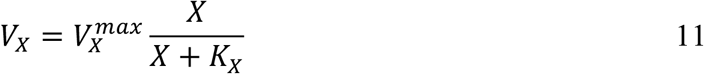

For dissolved oxygen, because it is a small, non-polar molecule that does not require active, enzyme-mediated transport into cells, we assume that the uptake rate 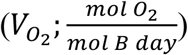 is limited by diffusive supply (*32*), which is dependent on the radius of the cell (r, metres), the diffusion coefficient of oxygen in seawater (*D*_*diff*_, m^2^ day^-1^), the cellular quota of nitrogen (Q_N_, mol N_bio_ cell^-1^) and the ambient oxygen concentration (O_2_, mol m^-3^) as:

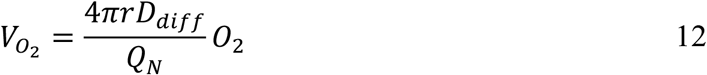

The radius of the cell is calculated from the cellular volume (Table S1), the diffusion coefficient is 1.5613 . 10^−5^ cm^2^ s^-1^, assuming 12°C and 35 psu (https://unisense.com/wp-content/uploads/2021/10/Seawater-Gases-table.pdf), and the cellular quota of nitrogen is calculated from estimates of carbon quotas and cell volume.

State variables are organic nitrogen (*N*_*org*_ or *OM* for yield notation), inorganic nutrients 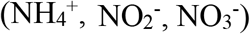, biologically-produced dissolved dinitrogen (N_2_) (“excess N_2_”, neglecting gas exchange with air), dissolved oxygen (O_2_) and the biomass of each microbial group (*B*_*NAR*_, *B*_*NIR*_, B_AOA_, B_NOB_, B_AOX_). Complete equations for each state variable in the chemostat are as follows:

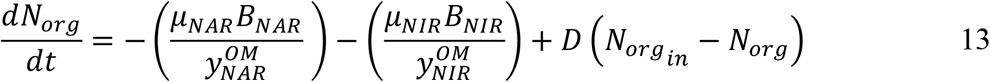

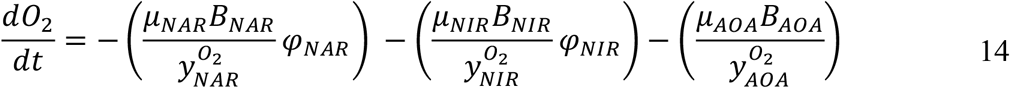

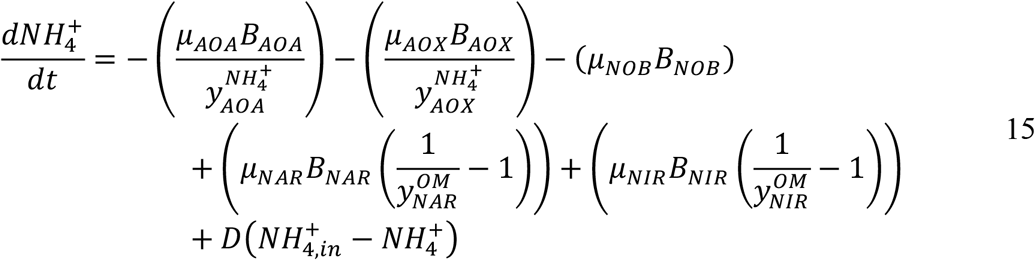

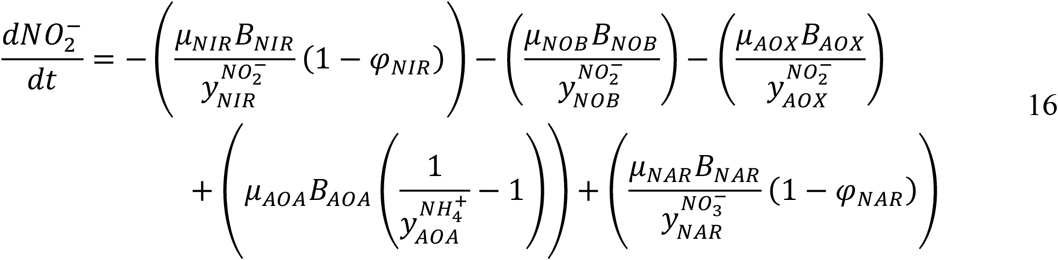

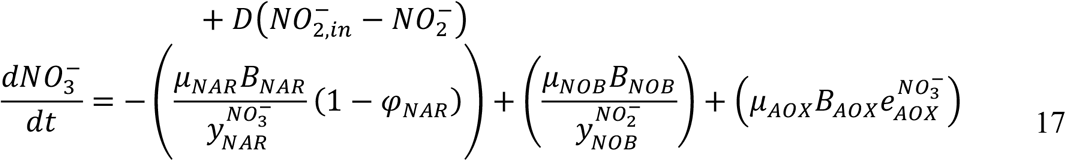

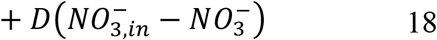

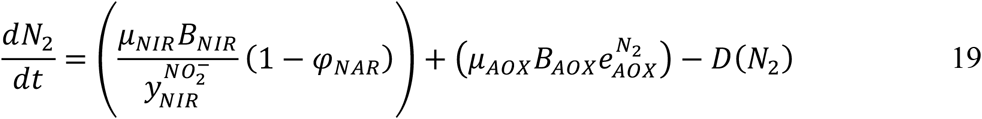

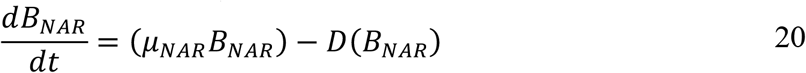

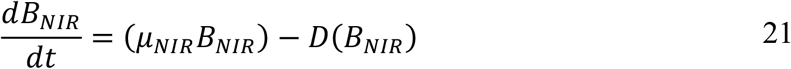

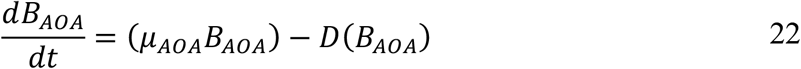

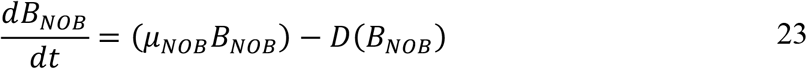

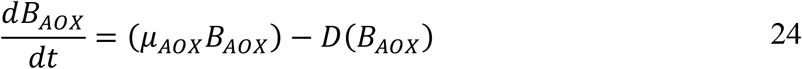

where *D* is the dilution rate (0.05 day^-1^), which sets the loss rate *L* and thus also the steady-state growth rate of the microbial populations in the chemostat.

Facultative heterotrophy (i.e., the switch between anaerobic and aerobic metabolism) was controlled with a Heaviside function stored in *φ*_*NAR*_ and *φ*_*NIR*_ for both the nitrate-reducing (NAR) and nitrite-reducing (NIR) heterotrophs as:

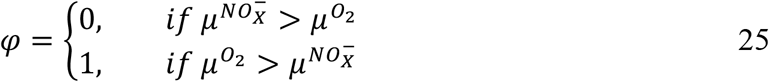

Equation 25 states that if growth rates achieved via aerobic metabolism are greater than the growth rates achieved by anaerobic metabolism (either nitrate or nitrite reduction), then the heterotroph will execute aerobic growth, and *vice versa*. Above, the *φ* term sets heterotrophic growth as either aerobic or anaerobic.

The default values of the yields of each metabolism are listed in Table S2.

#### Chemostat oxygen pulse experiments

We conducted pulsing experiments where oxygen was introduced to the chemostat model at a range of amplitudes and frequencies. Simulations were run for 10,000 days to achieve dynamic equilibrium (Fig. S3). These pulses were applied on top of constant, background rates of oxygen (0, 0.025 and 0.05 µM day^-1^) and organic matter (0.05 µM day^-1^) supply, which generated steady-state anoxia when pulses were not applied and excluded aerobic nitrifiers (red stars in Fig. 1). Oxygen pulses were instantaneous additions of oxygen of between 10 nM to 1 µM that occurred during a distinct timestep. Importantly, these pulses reoccurred over the simulation at a range of frequencies, from five times per day (0.2 day period) to once every 10 days. Every combination of pulse amplitude and period was explored, with extremes being 10 nM once every 10 days (e.g., AMZ core) and 1 µM every 0.2 days (e.g., oxygenated boundary). We therefore explored a landscape of pulse periods and amplitudes. In parallel experiments, we also heavily reduced the maximum growth rate of NOB from 1 day^-1^ to 0.5 day^-1^ (equal to heterotrophs) and 0.25 day^-1^ (equal to anammox) to test if our results were contingent on NOB’s rapid growth (Table S1). These experiments show no qualitative difference with those presented in the main text and indicate that while higher maximum growth rates of NOB increases the rate of their biomass accumulation during a “bloom” associated with an oxygen pulse, their higher maximum growth rates are not necessary for their survival, which is instead ultimately a function of their metabolic niche (Fig. S4).

#### Three-dimensional ocean-biogeochemical model

The biogeochemical model is the Biogeochemical Elemental Cycling (BEC) model (*40*), which has recently been updated to include a more complete description of nitrogen cycling (*72*). This biogeochemical model is coupled online to the Regional Ocean Modelling System (ROMS) (*73, 74*), which serves as the physical environment coupled to the BEC. Our southeast Pacific domain extends from 111.38ºW to 66.62ºW and from 42.52ºS to 3.41ºN. The horizontal resolution is nominally 1/10^th^ of a degree (10 km) and the vertical grid consists of 42 bathymetry-following levels with highest resolution at the surface and bottom. The same configuration of ROMS-BEC was recently used to explore N_2_O dynamics in the southeast Tropical Pacific (*41*), and more information regarding the physical and biogeochemical model configuration can be found there, including the repeat atmospheric forcing, initial conditions, and boundary inputs.

Biomass pools of facultative nitrate- and nitrite-reducing heterotrophs (NAR and NIR), anammox bacteria (AOX), ammonia-oxidizing archaea (AOA) and nitrite-oxidizing bacteria (NOB) were added to ROMS-BEC with the same traits and yields as in the chemostat model (Tables S1 and S2), although the values of the default maximum growth rates were adjusted so that they aligned with the maximum growth rates at 15 ºC as detailed below (Table S5). Furthermore, the implementation of explicit biomasses necessitated the addition of a nano-zooplankton functional type that exclusively grazed on the heterotrophic bacterial types. The chemoautotrophs, being on average larger than heterotrophic bacterial types (Table S1), were grazed by the zooplankton functional type already present within BEC. This size-based segregation of grazing reproduced the vertical patterns of abundance in heterotrophic bacteria and chemoautotrophs in the open ocean, whereby maximal heterotrophic biomass is observed in the euphotic zone, while maximal biomass of chemoautotrophs is observed just beneath the euphotic zone due to competition with phytoplankton for inorganic nutrients (*75, 76*).

Potential growth and grazing rates were made a function of temperature *T* in Kelvin as:

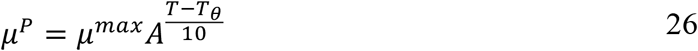

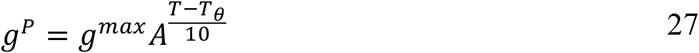

In the above, *A* is a constant and equal to 1.7, while maximum growth (*μ*^*max*^) or grazing (*g*^*max*^) rates are unique to the microbial or zooplankton functional type. To accommodate a reference temperature of 30 ºC (*T*_*θ*_ = 303.15 *K*), we doubled *μ*^*max*^ for all microbial functional types compared to the chemostat model, such that growth rates at ∼15 ºC were roughly 0.5, 0.5, 1.0 and 0.25 day^-1^ for heterotrophs, AOA, NOB and anammox bacteria, respectively, making them similar to those of the chemostat under these conditions (Table S1). Maximum grazing rates for the nano-zooplankton 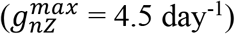 were faster 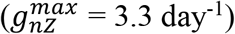 than the larger zooplankton type consistent with observations (*77*). Potential grazing rates of the larger zooplankton were also made a function of oxygen, such that grazing approached zero as oxygen approached zero as:

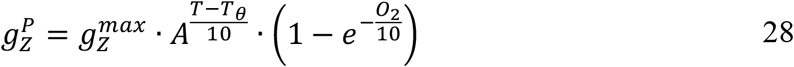

The deceleration of grazing in low oxygen zones was essential for ensuring a viable population of anammox bacteria, whose slow growth rates make them susceptible to extirpation by an active grazer.

Potential growth and grazing rates are then multiplied by limitation terms (*l*) dependent on resource availability to return realized growth and grazing rates, as:

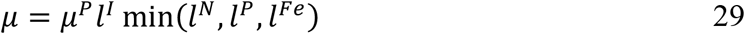

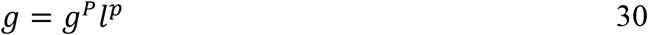

where the limitation terms are as follows:

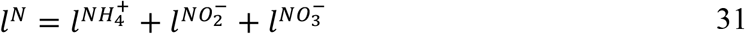

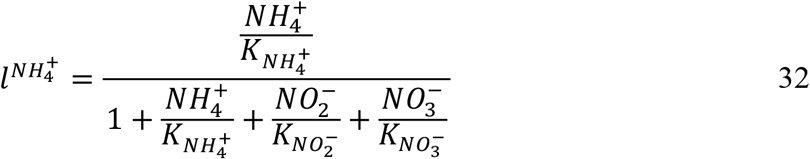

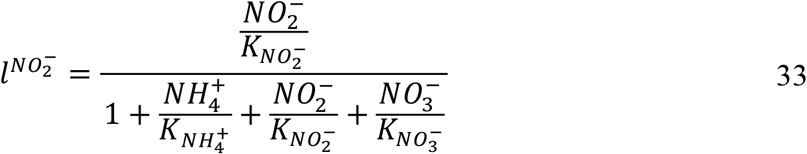

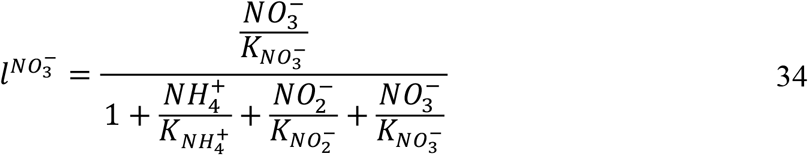

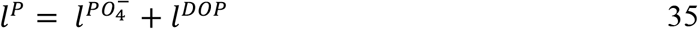

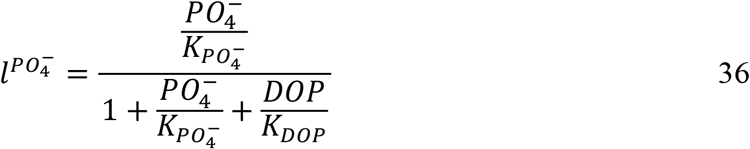

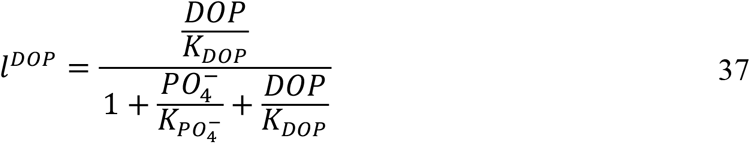

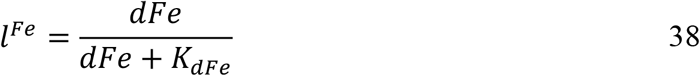

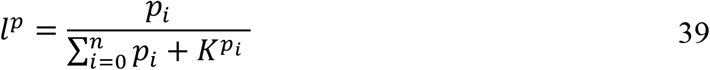

where DOP is dissolved organic phosphorus, dFe is dissolved iron, and *l*^*I*^ is light limitation of photosynthesis (described in Moore et al. (*40*)). Half-saturation coefficients and other parameters governing phytoplankton are provided in Table S8. For grazing on prey type *p*_*i*_, the model also takes the Monod form but normalizes grazing pressure by the total prey available 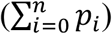, with 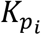 being the half-saturation coefficient specific to *p*_*i*_. For the prey of nano-zooplankton, being the heterotrophic microbial bacterial types (NAR and NIR), 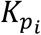 was greater than for the prey of the larger zooplankton type, being phytoplankton and chemoautotrophs, in keeping with observational constraints (*77*).

Although different stoichiometries of these microbial types are reported (Table S1), we chose to apply the same C:P, C:N and Fe:C stoichiometry of the heterotrophic and chemoautotrophic functional types for simplicity. These were 55:1 mol mol^-1^, 5:1 mol mol^-1^, and 20 µmol mol^-1^, respectively. Sensitivity experiments with different stoichiometries of C:P and C:N showed little effect. Heterotrophic bacteria obtain their carbon, nitrogen and phosphorus from the dissolved organic matter that they consume but assimilate dissolved bioavailable iron from the environment. All microbes obtained their iron this way, ensuring direct competition between photoautotrophs, chemoautotrophs and heterotrophs for iron, as observed (*78, 79*). We acknowledge that Fe:C quotas for these microbes can be much higher than 20 µmol mol^-1^ we ascribe here (*79*). Also, although phosphorus and iron are active prognostic cycles in the BEC model, we do not ascribe limitation terms to the growth of the microbial functional types based on the availability of these nutrients. These considerations will be explored in a future study.

All microbial functional types were prescribed the same linear and quadratic mortality rates, being 5% of their maximum growth rates and 0.2 (µM biomass day)^-1^, respectively. Linear mortality implicitly represents losses of biomass associated with cell respiration and maintenance costs, while quadratic mortality implicitly represents losses associated with viral lysis. Other considerations were the fraction of mortality and grazed biomass that was routed to grazer biomass, particulate organic matter, dissolved organic matter (both semi-labile and semi-recalcitrant types), and inorganic nutrients. For these parameters we reverted to the fractions already present for the existing ecosystem components in BEC, and thus conserved the behavior of the original model to the greatest possible extent. These and other parameters as input to the ocean-biogeochemical model are detailed in Tables S5 and S8.

We ran the ROMS-BEC model forward from initial conditions (see McCoy et al. (*41*)) for 51 years. Each timestep was 800 seconds. We present the results from the final year.

We also conducted three sensitivity experiments where the µ^max^ of the heterotrophic bacteria, AOA and NOB were altered sequentially (Table S4, S5) (Supplementary Text). First, heterotrophic bacteria µ^max^ was doubled from 1 day^-1^ to 2 day^-1^ at 30ºC; then the µ^max^ of NOB was halved from 2 day^-1^ to 1 day^-1^ at 30ºC; and finally, the third experiment doubled the µ^max^ of AOA from 1 day^-1^ to 2 day^-1^. Each experiment was branched from the control run at year 40 and run for 10 years in parallel.

#### Carbon quotas and yields for AOA and NOB

As in Zakem et al. (*44*), we convert modelled biomass to cell abundances using measured carbon quotas for AOA and NOB grown in natural seawater at 15°C and 1 µM substrate (*80*). For AOA, we use the average carbon content from *Ca. Nitrosopelagicus* U25 and *Nitrosopumilus* sp. CCS1: 11.5 ± 2.0 fgC per cell. For NOB, we use the carbon content of *Nitrospina* sp. Nb-3: 39.8 ± 11.2 fg C per cell. We also use the measured carbon fixation yields from these organisms as the biomass yields for AOA and NOB in our model (Table S1).

#### Estimating maximum growth rate from draft metagenomes

Metagenome-assembled genomes (MAGs) were generated from water samples taken in the anoxic zone in the Eastern Tropical South Pacific Ocean (*15*). We annotated each MAG using *prokka* (*81*) (with options --norrna --notrna --centre X --compliant and --kingdom Archaea when dealing with archaeal MAGs) to obtain coding sequences and locate ribosomal proteins. We then performed growth prediction as a function of codon usage bias of the ribosomal proteins with gRodon (*50*) assuming an optimal growth temperature of 13°C for all MAGs based on the mean of sample temperatures and using metagenome mode v2 (*82*) to control for any possible contamination caused by binning. Weissman et al. (*82*) provide a codon usage bias cut-off to classify an organism as a copiotroph or oligotroph (i.e., opportunists or gleaners (*39*). The cut off is based on where the codon usage bias versus minimum doubling time curve saturates (*50*). All predictions were done with gRodon’s new `training=“AOA_NOB”` flag that incorporates the new AOA and NOB training data (Table S7). This new model was trained using an identical procedure to the original gRodon model, with the addition of ten new NOB and AOA genomes and associated growth rates estimates from laboratory experiments.

#### Supplementary Text

##### Sensitivity experiments with the chemostat model

The traits detailed in Table S1 were used for predicting the outcome of competition for resources between microbes (resource-ratio theory) and as input parameters for both the chemostat and 3D ocean-biogeochemical models. Similarly, the yields of heterotrophic bacteria are known to vary over a wide range (*83*). However, some of these traits are not well constrained or are known to range widely, warranting a sensitivity analysis using *a priori* reasonable ranges.

We altered maximum growth rates, half-saturation coefficients for nutrient uptake, biomass yields, cellular carbon quotas and mortality terms to determine how sensitive the outcomes of the chemostat model were to these parameters. A qualitative difference was defined as a different combination of active microbial types under the same oxygen and organic matter supply ratio. For example, if nitrification replaced anaerobic heterotrophy under oxygen-limiting conditions, then a qualitative shift occurred.

Our model heterotrophs were based on SAR11 bacteria given their abundance in AMZs (*7, 18, 84*) and contribution to denitrification (*85*). In the real ocean, a diversity of microbes leads to a spectrum of O_2_*, known as the subsistence oxygen concentration on which an organism’s biomass gains equal biomass losses (see below for derivation of O_2_*). Accounting for a wide range of heterotrophic yields from 0.05 to 0.6 mol C_bio_ (mol C_org_)^-1^ (*83*) predicts multiple values of O_2_* ranging over two orders of magnitude. This indicates that some portion of the heterotrophic bacterial population may have higher O_2_* than AOA, but likely not higher than the O_2_* of NOB (Table S3) because of the latter’s much lower yield and larger cell size. This tells us that AOA may be able to competitive exclude aerobic heterotrophy in AMZs, or in other words, reduce the oxygen to a concentration that anaerobic, rather than aerobic, heterotrophy becomes the preferable metabolism for facultative heterotrophs. It seems unlikely, though, that NOB may contribute to the exclusion of a portion of aerobic heterotrophic metabolism. A heterotrophic biomass yield of roughly 0.06 mol C_bio_ (mol C_org_)^-1^ is required for AOA to outcompete heterotrophs for oxygen. This value sits well outside the average estimate for marine heterotrophy of 0.33 ± 0.1 (*86*). Furthermore, we see that the lower range of O_2_* for heterotrophic bacteria (0.15 nM) clearly falls beneath the estimated range for nitrifiers (AOA and NOB), suggesting that a portion of the heterotrophic community should always outcompete aerobic nitrifiers. Finally, the competitive upper-hand of heterotrophs is also robust to uncertainties in other parameters, including maximum growth rates, nutrient affinities, carbon quotas, as well as a constant versus quadratic mortality rate (Fig. S1). It is noteworthy that our results were not changed when heterotrophic bacteria had higher growth rates than NOB.

##### Competitive outcomes of the chemostat model

The competitive outcomes of the chemostat model can be anticipated by solving for the population-specific resource subsistence concentrations, R* (*31*), by assuming steady-state such that growth (*µ*) equals losses (L) of biomass. For organic matter and inorganic nutrients, the R* is calculated for resource *i* and population *j* as:

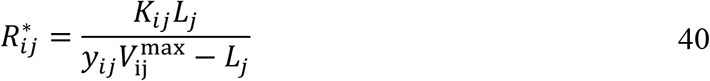

For dissolved oxygen, O_2_*, we use the equation derived in Zakem & Follows (*32*) for population *j*:

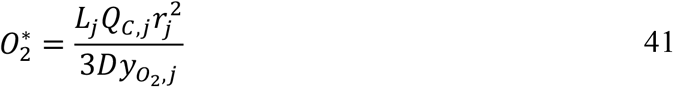

Where the carbon quota of the cell (Q_C_), radius (r), diffusion coefficient (D) and yield 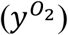 are in units of mol C µm^-3^, µm, µm^2^ day^-1^ and mol C_bio_ (mol O_2_)^-1^, respectively. This equation will return similar values to an R* derived from Michaelis-Menton kinetics (Eq. 40) when considering very high affinities (very low 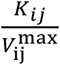).

##### Sensitivity experiments in ROMS-BEC

We calculate the contribution of nitrite oxidation to oxygen consumption in the modeled AMZ across four simulations in which we vary the maximum growth rate of the aerobic functional types (Fig. 4b, Fig. S9b): 1) where the *μ*^*max*^ of the NOB population is twice that of the facultatively aerobic heterotrophic bacterial populations (our default model); 2) where *μ*^*max*^ is the same for both NOB and heterotrophs, by increasing the *μ*^*max*^ of heterotrophs; 3) where *μ*^*max*^ of NOB is half that of heterotrophic bacteria; and 4) where *μ*^*max*^ of AOA was doubled to equal that of the heterotrophs. See Table S4 for more detail. For each individual simulation, the illustrated variation in the contribution reflects variations in time. Across these simulations, nitrite oxidation contributes 24-49% to oxygen consumption in waters where oxygen is less than 1 µM. Importantly, however, when the linear mortality constant was simultaneously adjusted (so that mortality scaled with maximum growth rate), there was very little difference in this estimated contribution (all converged to the statistics of the default model). This emphasizes how multiple parameters in the model that control the growth and loss rates of the microbial population may impact the model estimates. While all of these scenarios involve uncertainty, we think that the default model may best capture reality because it is known that the most abundant and ubiquitous bacteria in AMZs, as in the whole ocean, are members of the slow-growing SAR11 clade (*18, 48*). Fast-growing taxa are a smaller fraction of heterotrophic abundance, but we acknowledge that they may nonetheless contribute a disproportionate amount to the biomass that grows rapidly during an oxygen intrusion, and so may contribute more substantially to oxygen consumption than their relative abundance alone would predict (*87*). We therefore provide an upper bound on this scenario in our experiment where all heterotrophs can grow twice as fast as NOB. Though the resulting contribution to total oxygen consumption decreases to 24%-32%, this contribution is still substantial. Critically, these sensitivity experiments demonstrate the robustness of our main conclusions: that time-varying supply of oxygen is the mechanistic explanation for the observed high abundance of NOB and high rates of (aerobic) nitrite oxidation in AMZs. Future work can examine the impact when the diversity of the heterotrophic population is considered explicitly.

**Fig. S1.**
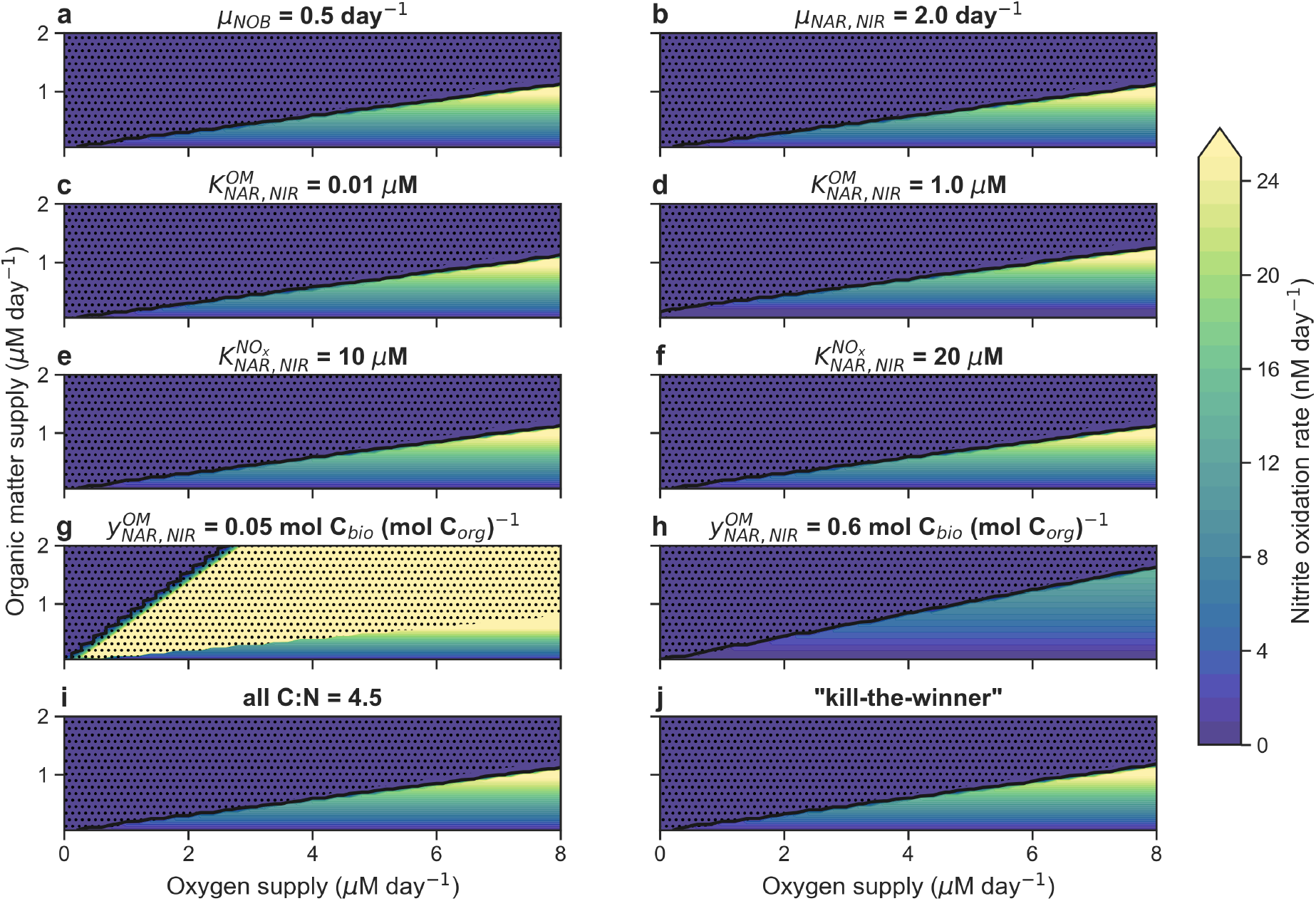
Sensitivity tests with chemostat model. Uncertain parameters of different traits are tested in each panel and run to equilibrium over 1640 combinations of oxygen and organic matter supply rates. (**a**) Growth rate of nitrite-oxidising bacteria halved from 1.0 to 0.5 day^-1^. (**b**) Growth rate of facultative heterotrophic bacteria quadrupled from 0.5 to 2.0 day^-1^. (**c**) Half-saturation coefficient for organic matter of facultative heterotrophic bacteria decreased an order of magnitude. (**d**) Half-saturation coefficient for organic matter of facultative heterotrophic bacteria increased an order of magnitude. (**e**) Half-saturation coefficient for nitrite and nitrate of facultative heterotrophic bacteria increased from 4 to 10 µM. (**f**) Half-saturation coefficient for nitrite and nitrate of facultative heterotrophic bacteria increased from 4 to 20 µM. (**g**) Yield of facultative heterotrophic bacteria decreased to 0.05 mol C_bio_ (mol C_org_)^-1^. (**h**) Yield of facultative heterotrophic bacteria increased to 0.6 mol C_bio_ (mol C_org_)^-1^. (**i**) All microbes have the same carbon to nitrogen ratio. (**j**) Mortality rate proportionally dependent on biomass in a ‘kill-the-winner’ approach. Black contour line is 1 nM oxygen concentration. Stippled region is where anaerobic heterotrophy is present.

**Fig. S2.**
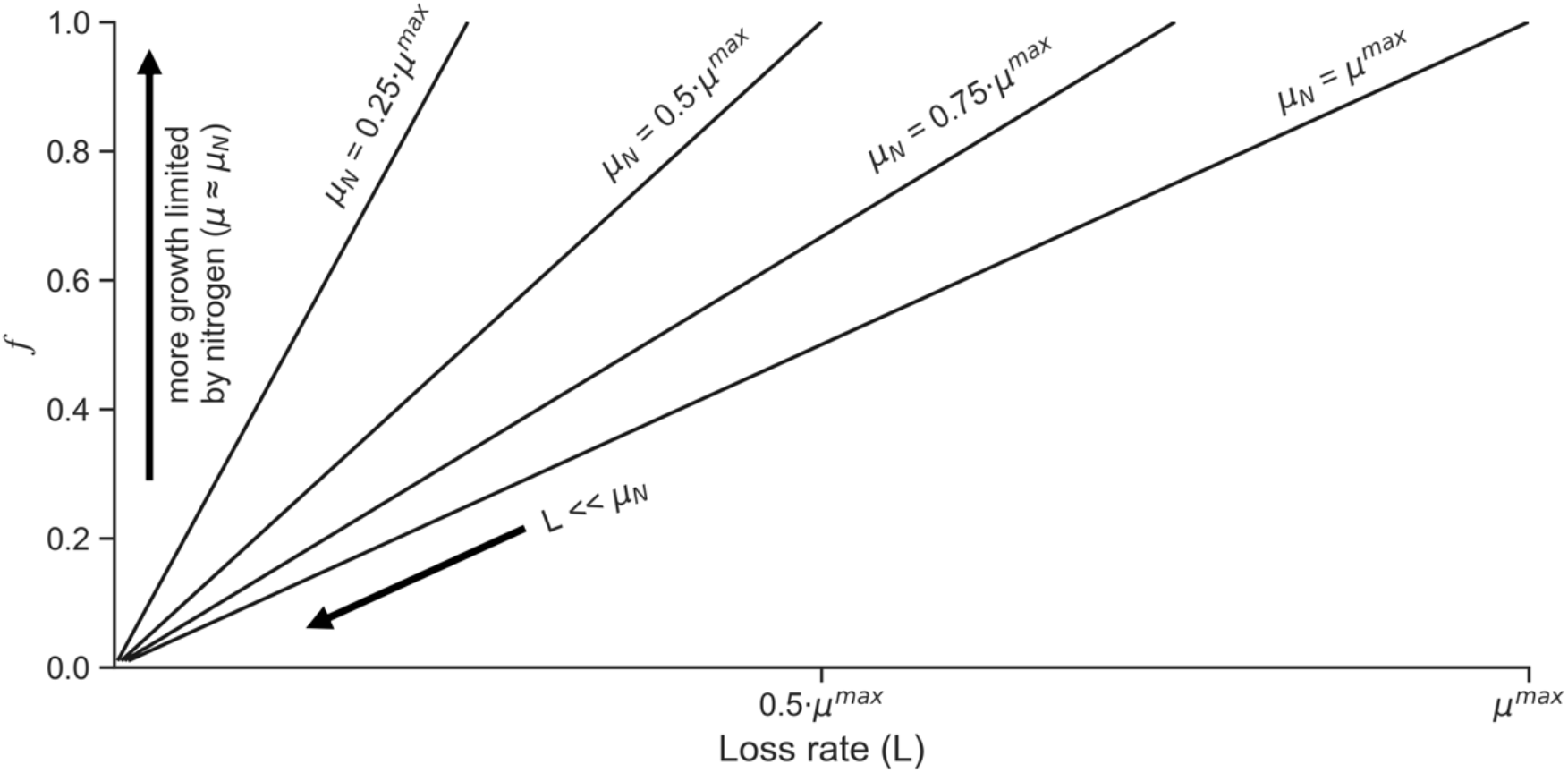
Relationship between *f, μ*_*N*_ and *L* for nitrifier growth in AMZs. The fraction of time, *f*, that nitrifiers require a switch from 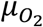 to *μ*_*N*_ to sustain their biomass is a function of how rapidly they can grow on nitrogen when released from oxygen limitation (*μ*_*N*_) and their loss rate (*L*) via Eq. 3. If *L* ≪ *μ*^*max*^, then *f* nears 0, and infrequent oxygen intrusions can sustain nitrifier biomass. However, if nitrogen is scarce in the environment and limits *μ*_*N*_, such that *μ*_*N*_ ≪ *μ*^*max*^, then nitrifiers need more oxygen intrusions (i.e., *f* increases) to sustain their biomass.

**Fig. S3.**
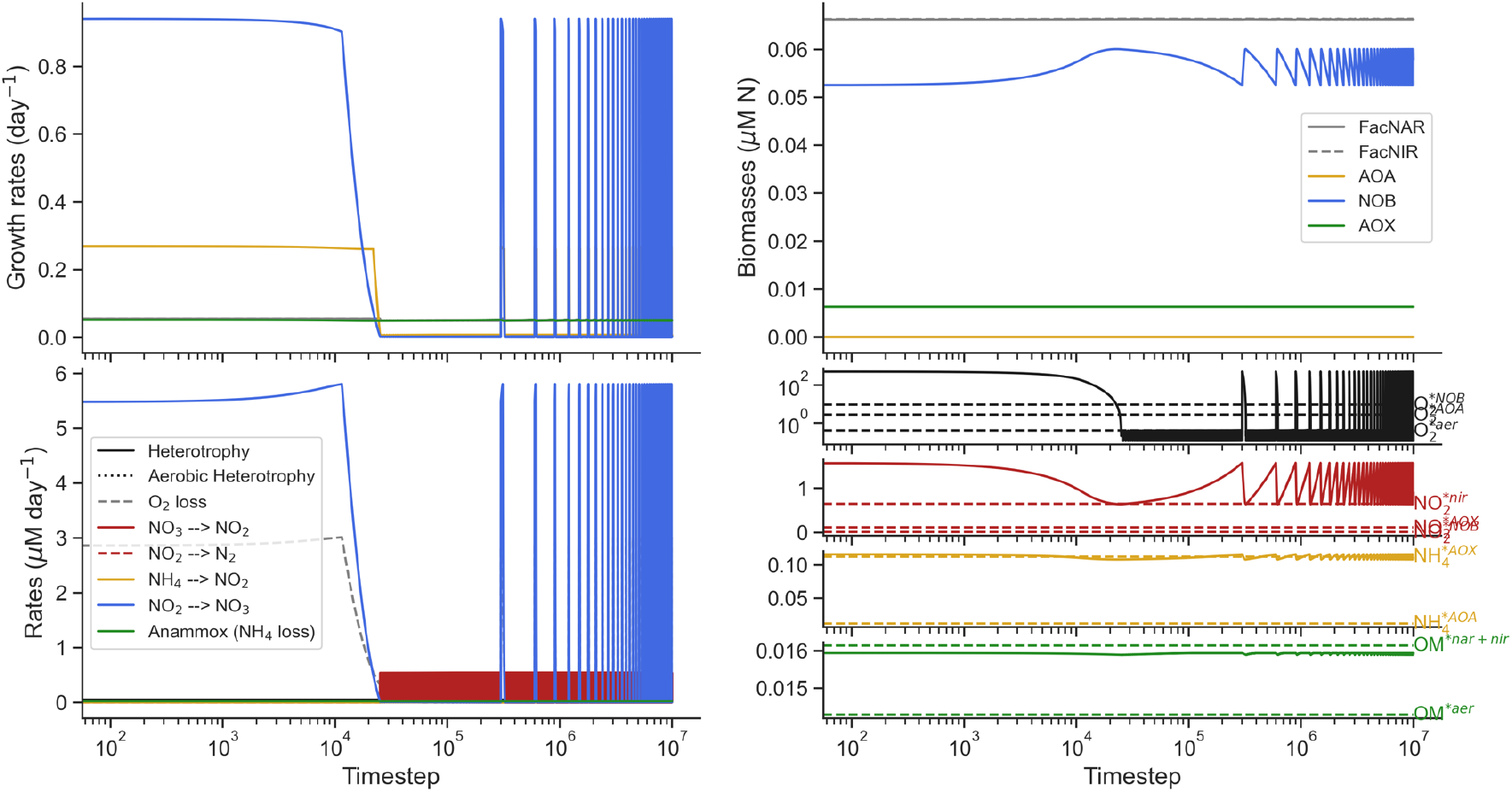
Example timeseries of pulse experiments. In this example, oxygen pulses are 1000 nM and occur every 6 days with a background constant oxygen supply of 0.025 µM day^-1^ and organic matter (N_org_) supply of 0.05 µM day^-1^ (red star in Fig. 1b,c). Growth rates (top left), rates of main metabolisms (bottom left), biomasses of microbial functional types (top right), and the concentrations of key tracers (bottom right), being oxygen (black), nitrite (red), ammonium (yellow) and organic matter (green). Oxygen (y-axis) and timesteps (x-axes) are shown on a log_10_ scale. Each timestep is 1000^th^ of one day, and the experiment is run for 10,000 days. The subsistence resource concentration required to sustain a viable population under steady-state conditions is indicated next to the plots of each tracer on the bottom right. “aer” = aerobic heterotrophy. “nar” = nitrate reducing denitrifiers. “nir” = nitrite reducing denitrifiers. “NOB” = nitrite oxidizing bacteria. “AOA” = ammonia oxidizing archaea. “AOX” = anammox bacteria. “FacNAR” = facultative nitrate-reducing heterotrophs. “FacNIR” = facultative nitrite-reducing heterotrophs.

**Fig. S4.**
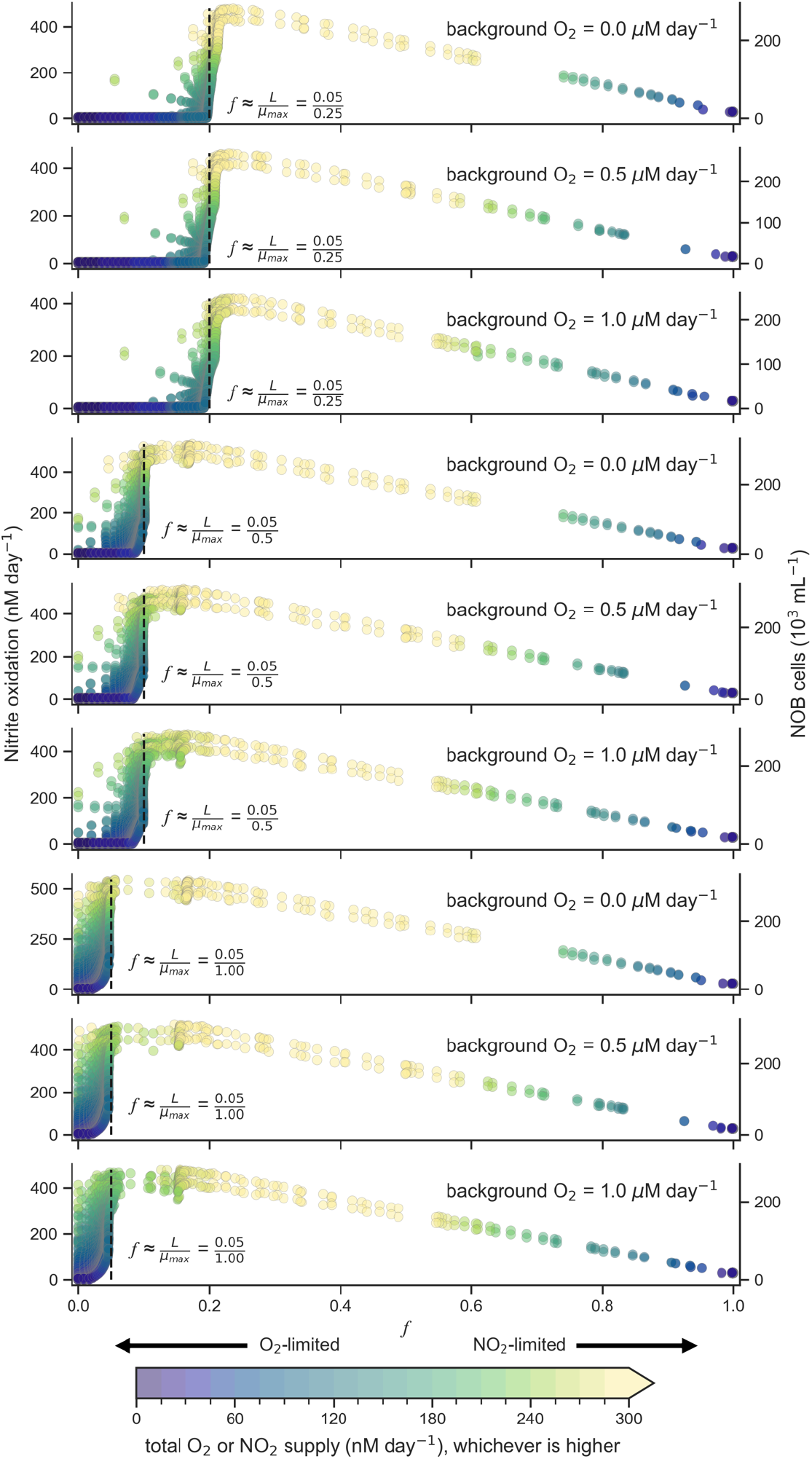
Effect of changing the maximum growth rate of NOB and background O_2_ supply rates. Nitrite oxidation rates are shown on the left-hand y-axis and NOB cells (1000s per mL) on the right-hand y-axis. X-axis is the model-derived *f* value, conceptually the same as the fraction of time NOB growth is released from O_2_ limitation. In each panel, from top to bottom, we have varied either the maximum growth rate of NOB and/or the underlying O_2_ supply rate. Note that *f* is sensitive to the chosen µ^max^ of NOB, but nonetheless the highest biomass and activity of NOB consistently approach the theoretically derived *f* value of 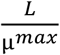. Varying underlying O_2_ supply has little effect.

**Fig. S5.**
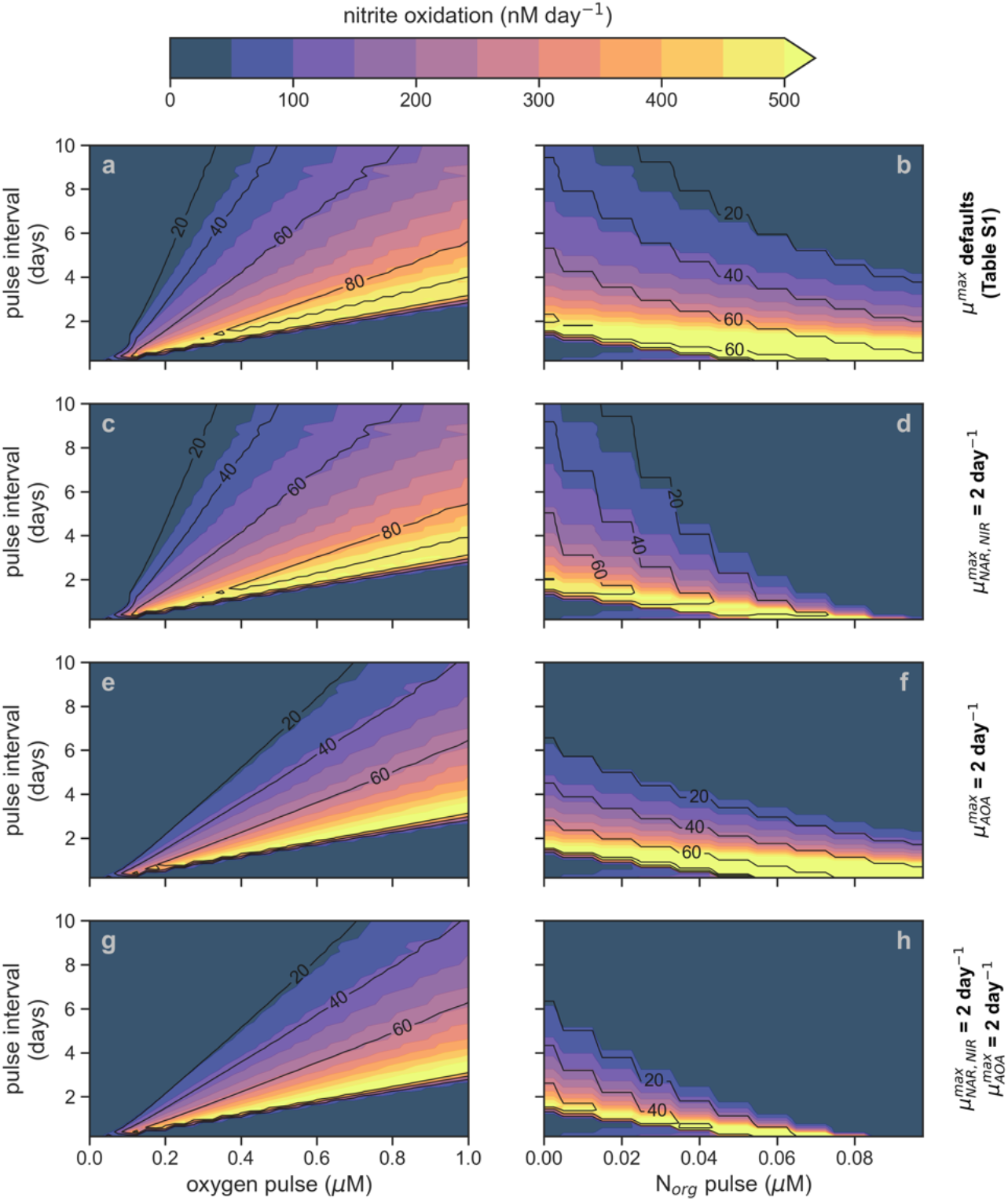
Nitrite oxidation rates (shading) and percent contribution of NOB to oxygen consumption (contours) from sensitivity experiments testing the effect of fast-growing (i.e., µ^max^ = 2.0 day^-1^) heterotrophs and AOA. NOB µ^max^ = 1 day^-1^. Oxygen pulsing experiments ranging from 0 to 1 µM at periods ranging from 0 to 10 days are shown on the left (panels **a, c, e**, and **g**). Organic matter (in form of nitrogen; N_org_) pulsing experiments ranging from 0 to 0.1 µM at periods ranging from 0 to 10 days are shown on the right (panels **b, d, f**, and **h**). Every experiment involves a constant background supply of both oxygen (at 0.025 µM day^-1^) and N_org_ (at 0.05 µM day^-1^) on top of which the pulsing of resources occurs (red star in Fig. 1b,c). The N_org_ pulsing experiments (right side) additionally include oxygen pulsing at an amplitude of 0.5 µM, such that a pulse of N_org_ is delivered alongside a pulse of 0.5 µM of oxygen. Hence, the nitrite oxidation rates at a N_org_ pulsing of 0.0 µM on the right are the same as the nitrite oxidation rates recorded at an oxygen pulse of 0.5 µM on the left. “NAR” = nitrate reducing denitrifiers. “NIR” = nitrite reducing denitrifiers. “AOA” = ammonia oxidizing archaea. “NOB” = nitrite oxidizing bacteria.

**Fig. S6.**
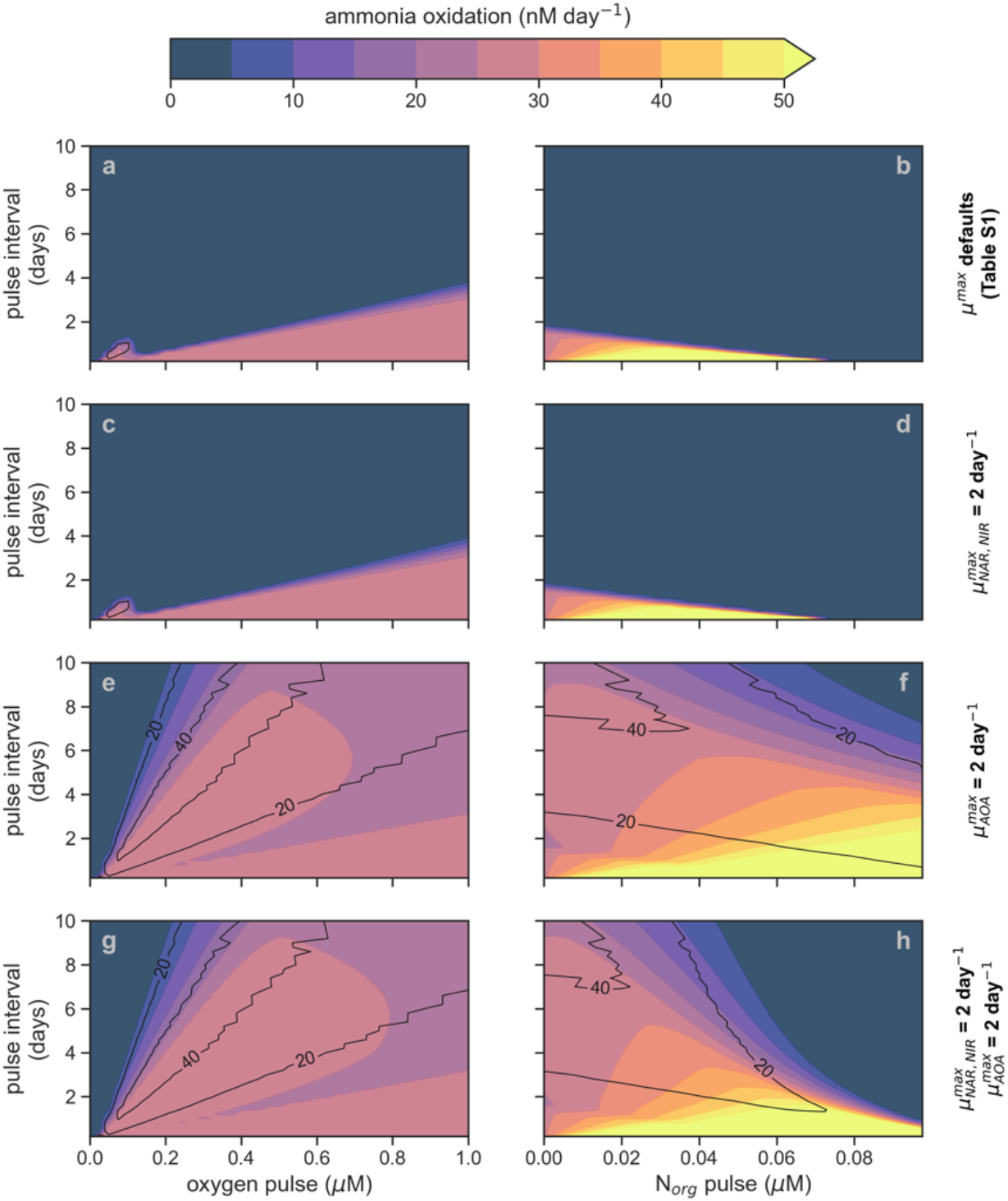
Ammonia oxidation rates (shading) and percent contribution of AOA to oxygen consumption (contours) from sensitivity experiments testing the effect of fast-growing (i.e., µ^max^ = 2.0 day^-1^) heterotrophs and AOA. NOB µ^max^ = 1 day^-1^. Oxygen pulsing experiments ranging from 0 to 1 µM at periods ranging from 0 to 10 days are shown on the left (panels **a, c, e**, and **g**). Organic matter (in form of nitrogen; N_org_) pulsing experiments ranging from 0 to 0.1 µM at periods ranging from 0 to 10 days are shown on the right (panels **b, d, f**, and **h**). Every experiment involves a constant background supply of both oxygen (at 0.025 µM day^-1^) and N_org_ (at 0.05 µM day^-1^) on top of which the pulsing of resources occurs (red star in Fig. 1b,c). The N_org_ pulsing experiments (right side) additionally include oxygen pulsing at an amplitude of 0.5 µM, such that a pulse of N_org_ is delivered alongside a pulse of 0.5 µM of oxygen. Hence, the nitrite oxidation rates at a N_org_ pulsing of 0.0 µM on the right are the same as the nitrite oxidation rates recorded at an oxygen pulse of 0.5 µM on the left. “NAR” = nitrate reducing denitrifiers. “NIR” = nitrite reducing denitrifiers. “AOA” = ammonia oxidizing archaea. “NOB” = nitrite oxidizing bacteria.

**Fig. S7.**
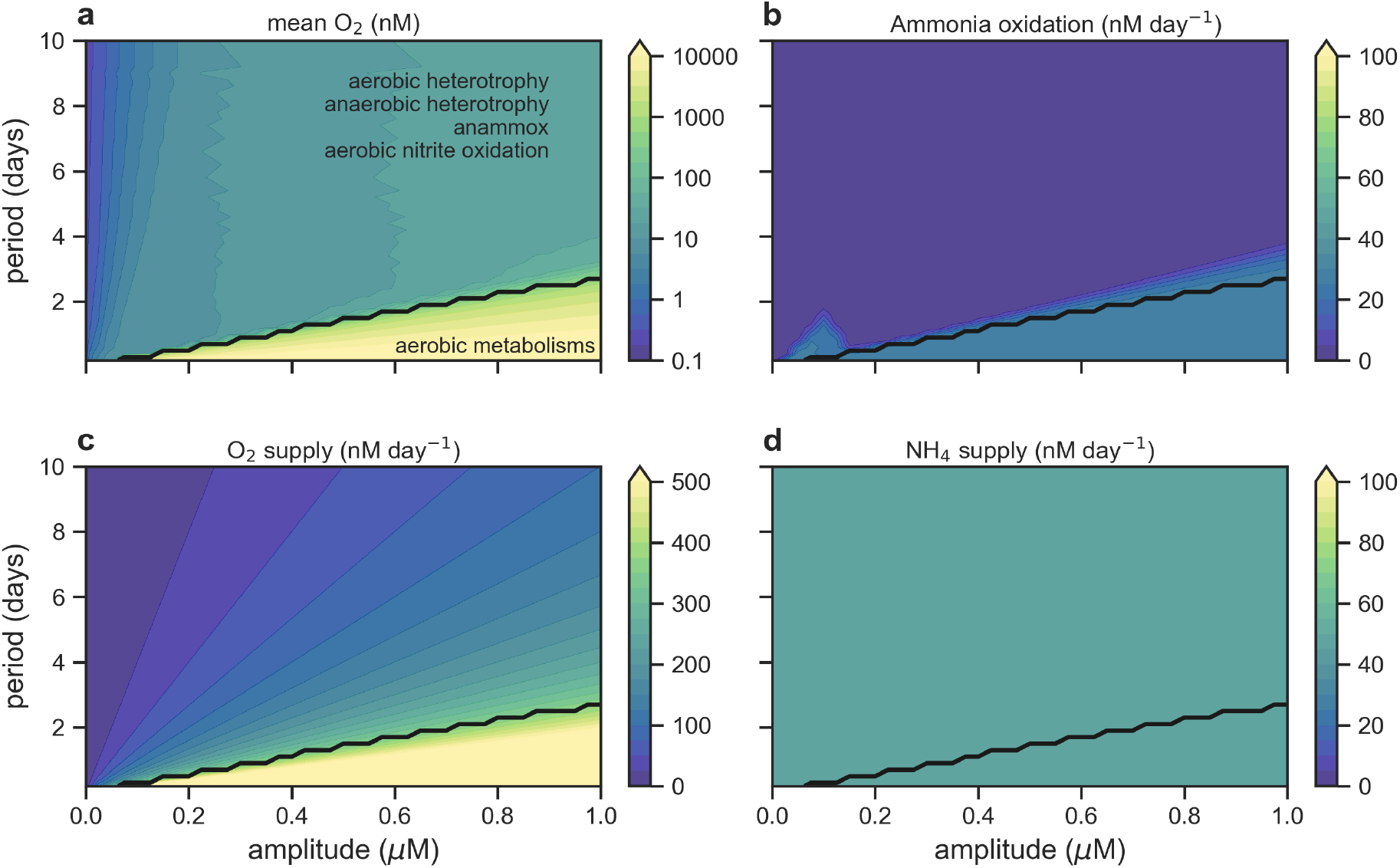
Episodic oxygen pulses applied to the chemostat model do not support ammonia oxidation. (**a**) Average concentration of dissolved oxygen (O_2_) in nanomolar (nM) over last 100 days of chemostat experiments (run to equilibrium) with background oxygen and organic matter supply of 0.5 and 1.0 µM day^-1^. Black line represents threshold where 100 % of heterotrophy becomes aerobic. (**b**) Average rate of ammonia oxidation in nM day^-1^. (**c**) Rate of oxygen supply (nM day-1). (**d**) Rate of ammonium supply (nM day^-1^).

**Fig. S8.**
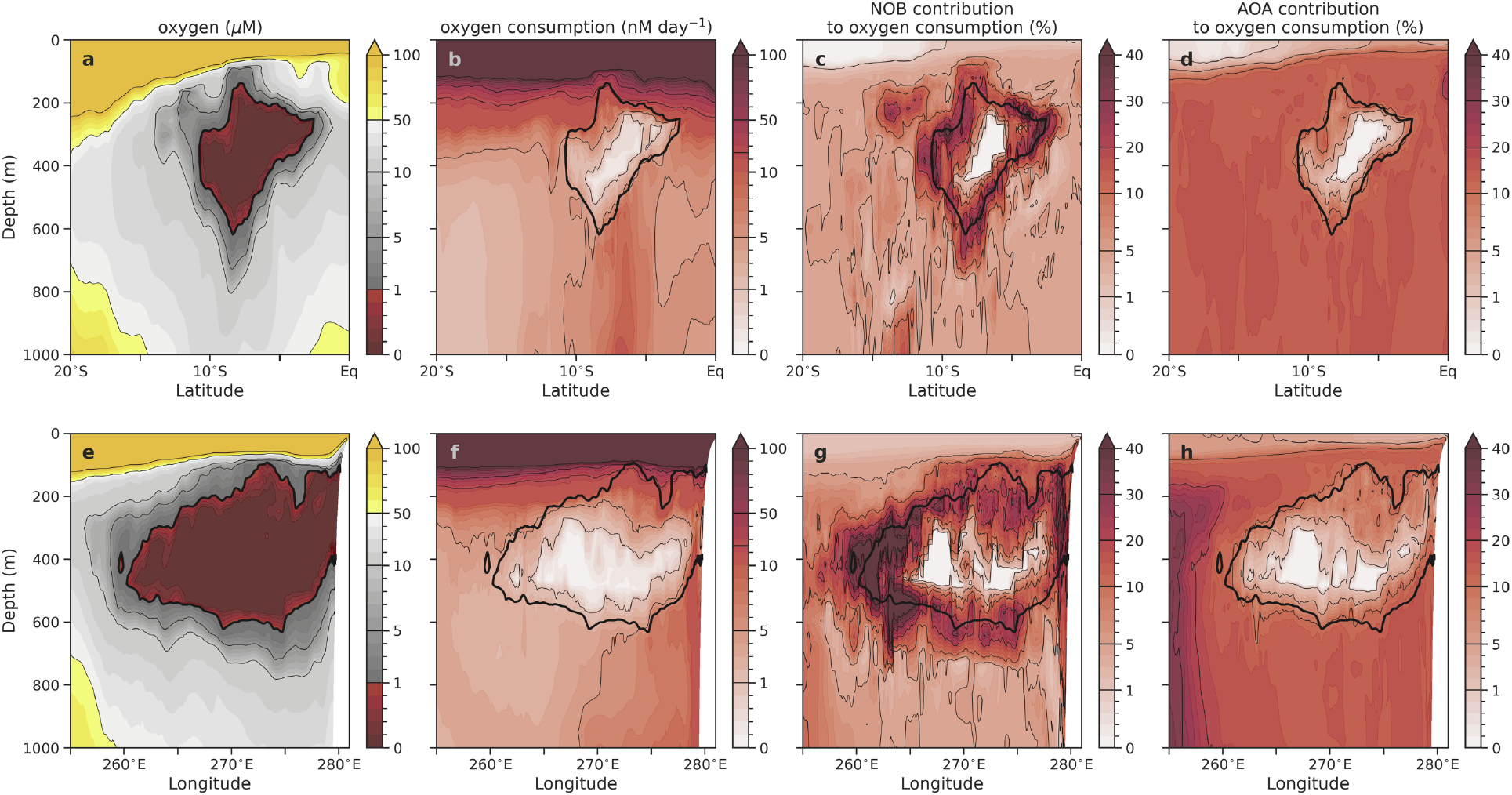
Contribution of NOB and AOA to total oxygen consumption. Top row is oxygen concentration (**a**), total oxygen consumption rate (**b**) and contribution of NOB (**c**) and AOA (**d**) to oxygen consumption along a latitudinal section through the AMZ at a longitude of 85ºW. Bottom row is the same but along a longitudinal section through the AMZ at a latitude of 8ºS. Values represent annual averages after 50 years of the control simulation from initial conditions. Heterotrophic contribution to oxygen consumption (not shown) is the remainder after accounting for NOB and AOA.

**Fig. S9.**
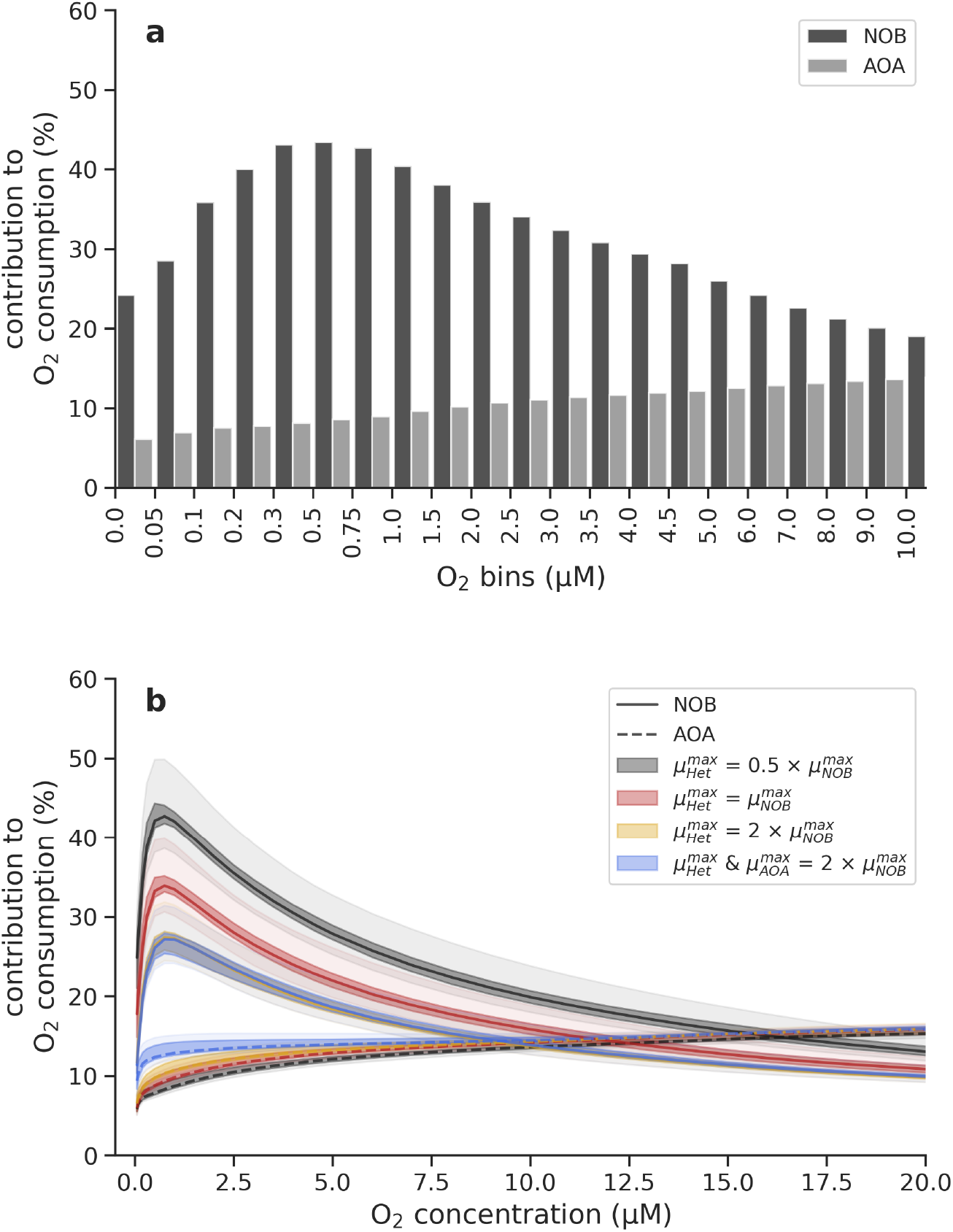
Integrated contributions of NOB (black) and AOA (grey) to total oxygen consumption, calculated cumulatively beneath discrete oxygen concentrations. (**a**) Annually averaged mean. (**b**) Annual median, interquartile (25^th^-75^th^ percentile) and 1^st^-99^th^ percentile ranges of the integrated contribution to O_2_ consumption for both NOB and AOA in the default model (black) and for sensitivity experiments that varied assumed maximum growth rates (µ^max^). Variations are in time. Data comes from daily mean output of the 51^st^ year of the 3D ocean-biogeochemical simulation.

**Fig. S10.**
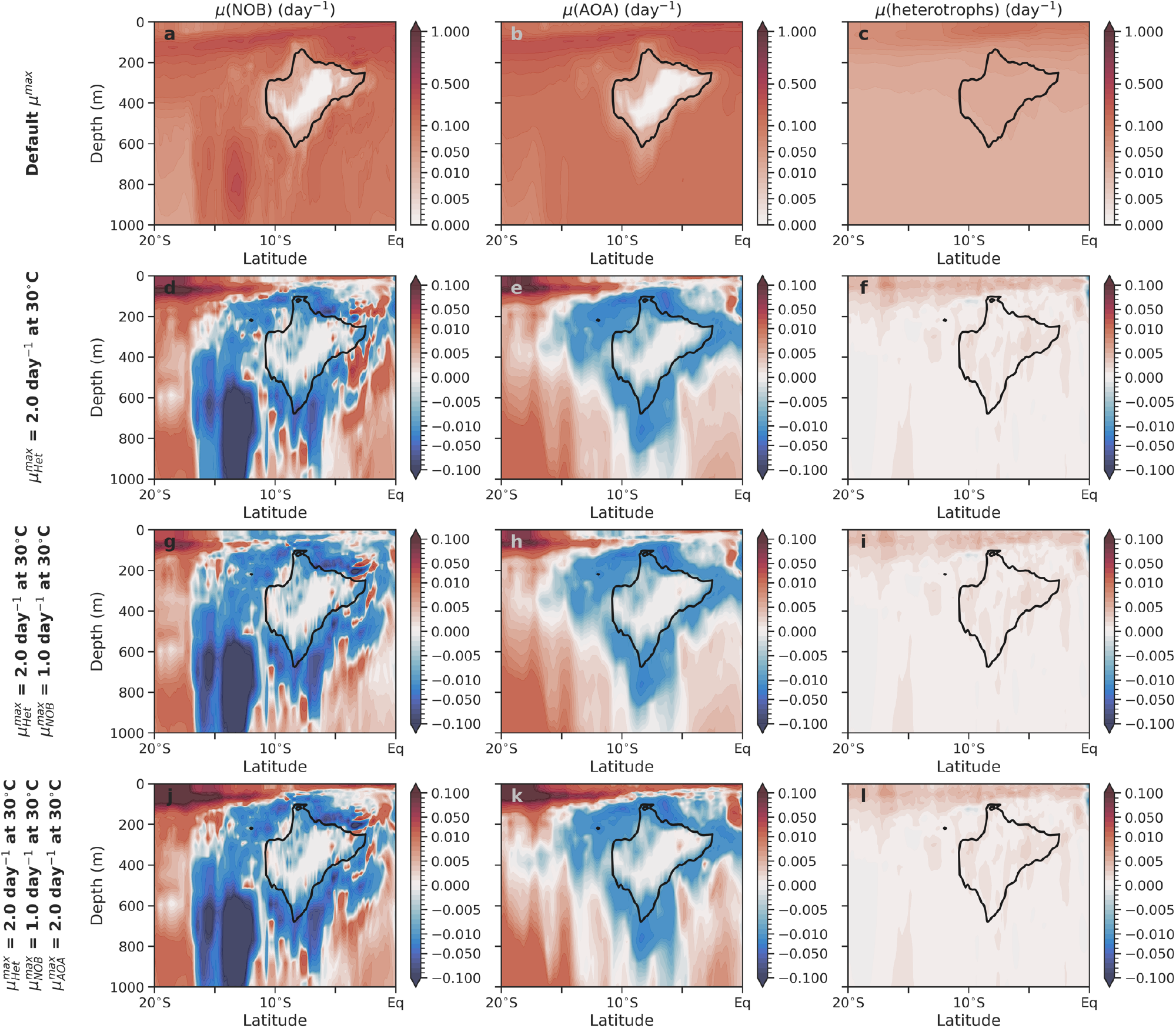
Realized growth rates of major microbial types and changes associated with the sensitivity experiments. Top row is µ of NOB (**a**), AOA (**b**) and facultative heterotrophic bacteria (**c**) along a latitudinal section through the AMZ at a longitude of 85ºW. Subsequent rows show changes in the same realized growth rates but for experiments with altered µ^max^ of one or more types (read left). Values represent annual averages after 50 years of the control simulation from initial conditions. Default µ^max^ values for the ocean-biogeochemical model are in Table S5.

**Fig. S11.**
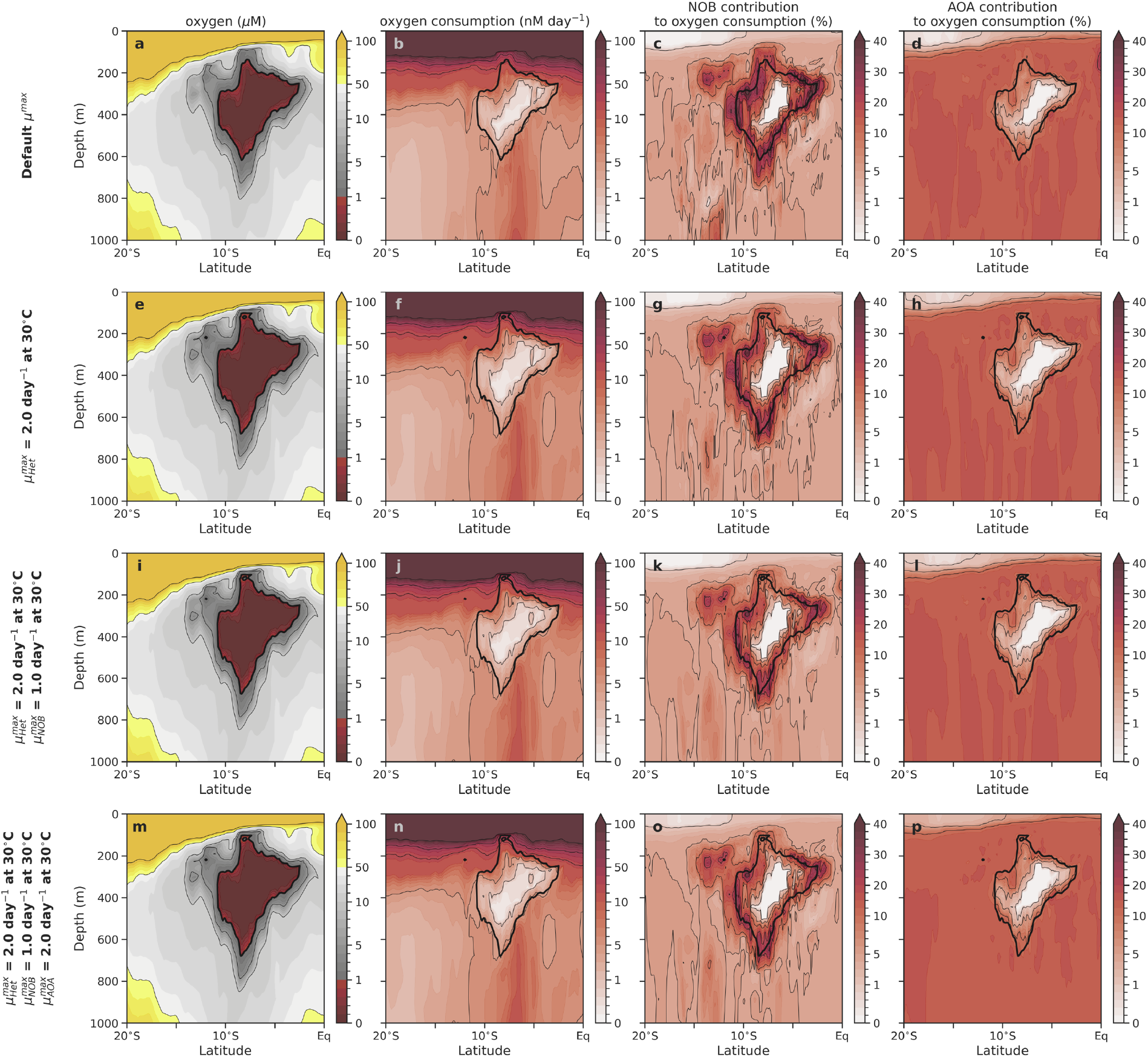
Contribution of NOB and AOA to oxygen consumption in sensitivity experiments. Oxygen concentration (**a**), total oxygen consumption rate (**b**) and contribution of NOB (**c**) and AOA (**d**) to oxygen consumption along a latitudinal section through the AMZ at a longitude of 85ºW. Subsequent rows show the same properties but for sensitivity experiments with altered µ^max^ of heterotrophs, NOB and/or AOA (read left). Values represent annual averages after 50 years of the control simulation from initial conditions. Heterotrophic contribution to oxygen consumption is the remainder after accounting for NOB and AOA. Default µ^max^ values for the ocean-biogeochemical model are in Table S5.

**Fig. S12.**
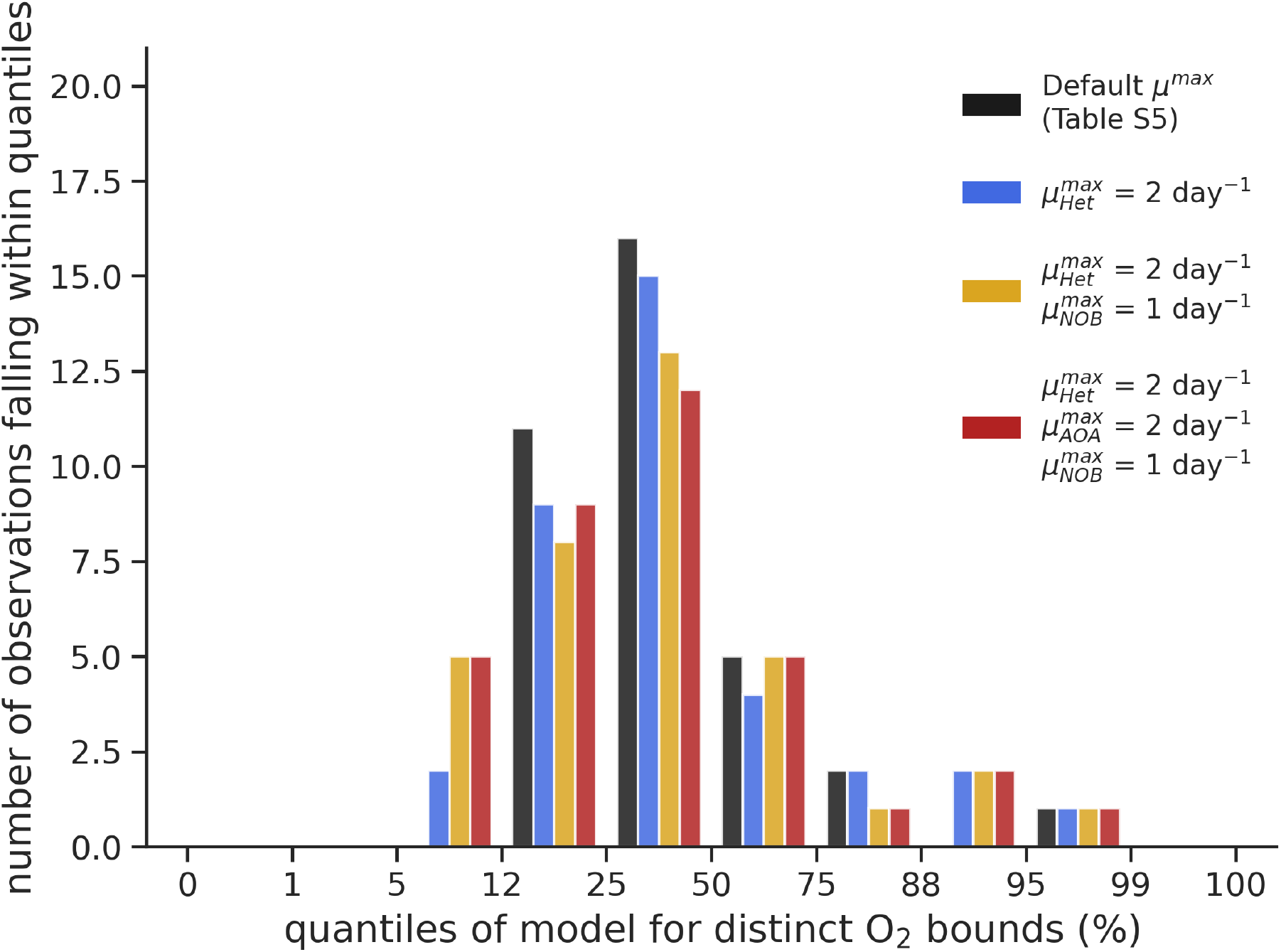
Frequency of occurrence of observed NOB contribution to oxygen consumption within the predicted frequencies from the model across sensitivity experiments. Model frequencies are calculated within discrete oxygen bounds and observations are allocated to a given model frequency range appropriate to their oxygen concentration. This analysis shows that the observed frequency of contributions matches well with that predicted by our model and that no experiment is better aligned with the observations than another. Figure made using monthly mean output over final year.

**Fig. S13.**
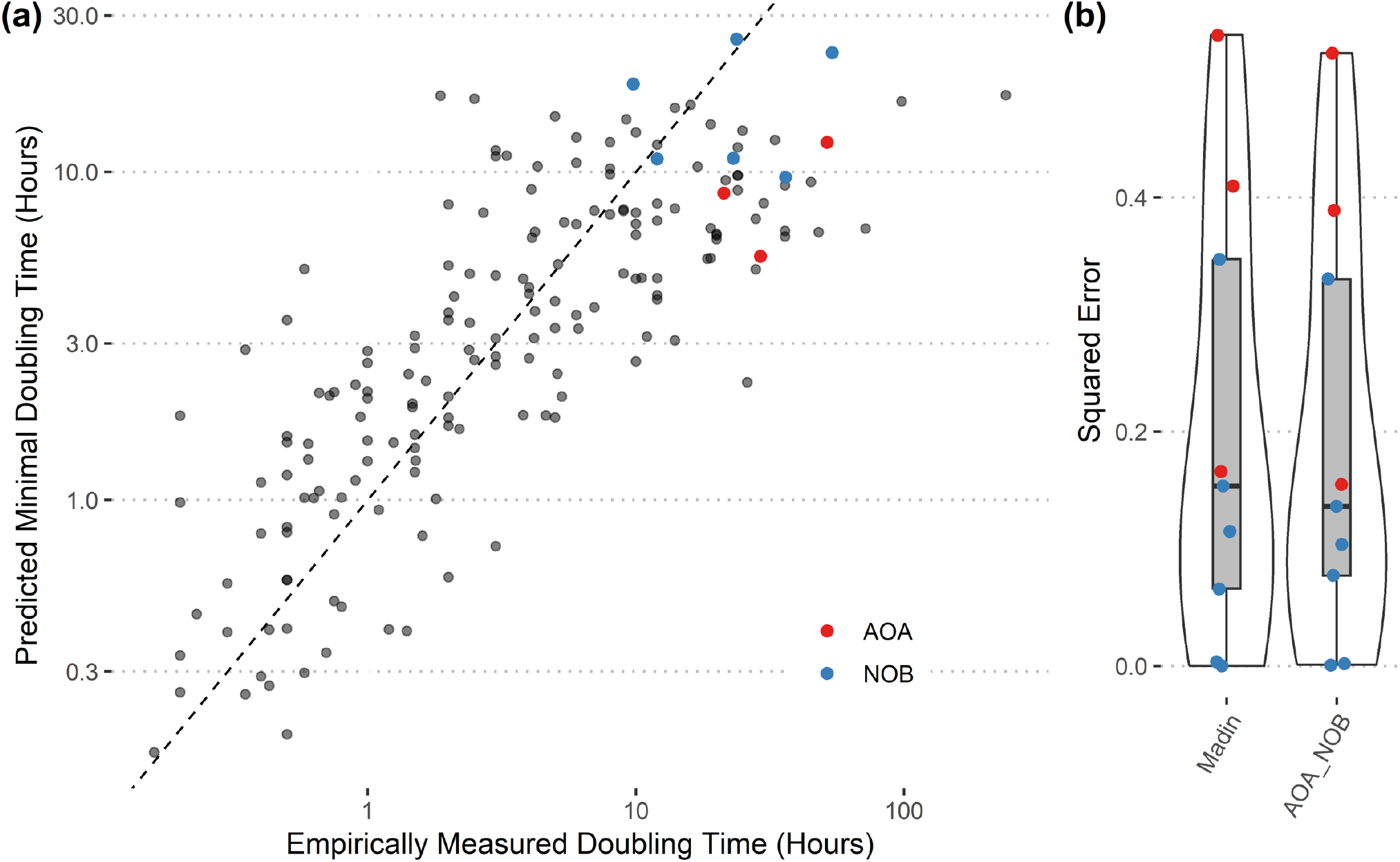
Performance of the updated gRodon statistical model for estimating maximum growth rates of key microbes. (**a**) Measured and predicted doubling times. Red dots are the new AOA and blue dots are the new NOB (chemoautotroph) data added to the model. (**b**) Square error of the predicted doubling times of the new chemoautotroph data using the previous (Madin) and updated (AOA_NOB) statistics.

**Fig. S14.**
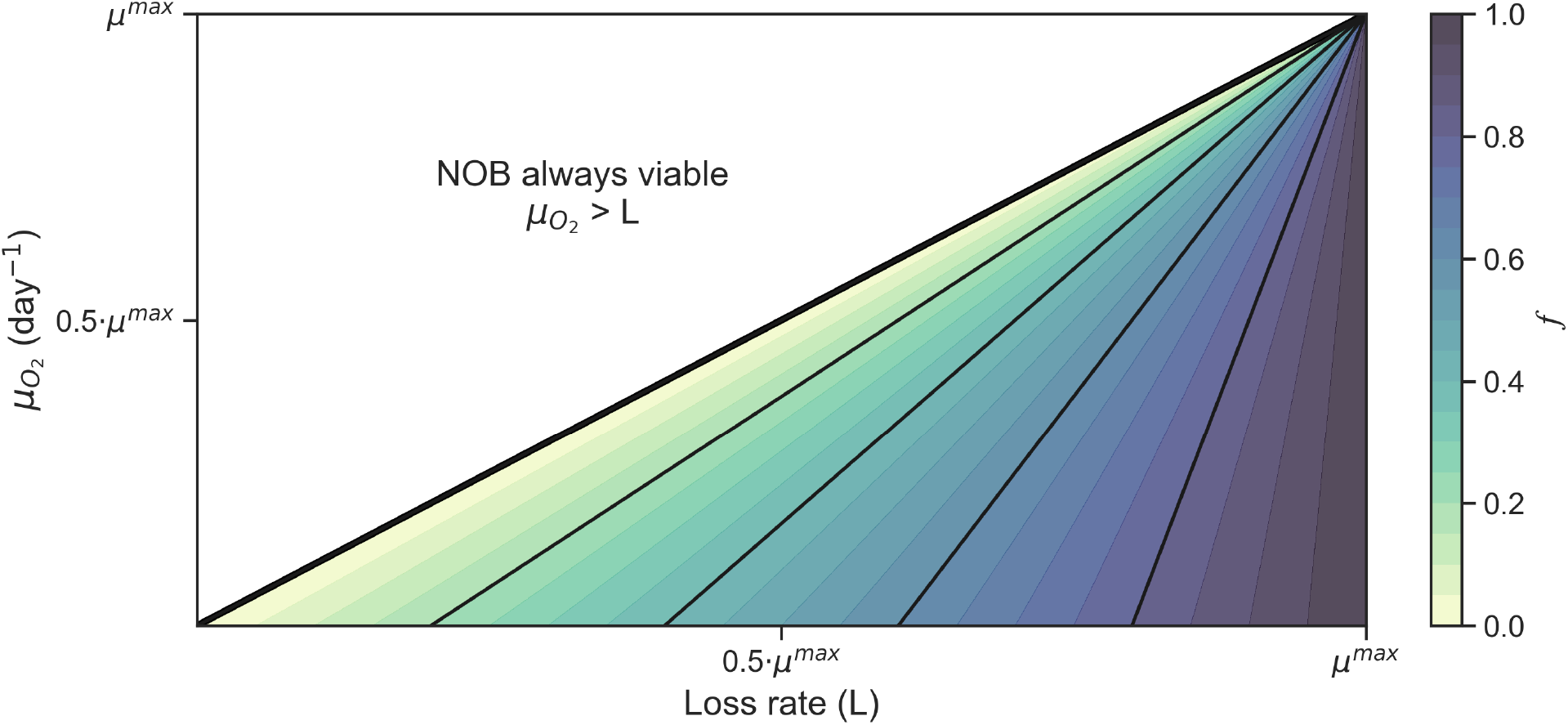
Quantitative demonstration of how anaerobic growth can make NOB more viable in low oxygen conditions. Here, ***f*** is a function of loss rate and growth under oxygen-limited conditions 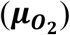 via Eqn. 2 in main article. In that equation, 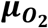 may be greater than zero and decreases ***f***. Nitrifiers capable of anaerobic growth require less frequent oxygen intrusions, or potentially none if anaerobic growth under oxygen limitation is greater than the loss rate 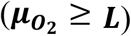.

**Table S1.** Empirically estimated traits of major marine microbial groups involved in key nitrogen transformations. These traits are assembled as input to the chemostat model and for an expanded analysis assessing the competitive outcomes of the different functional types when oxygen is limiting (Fig. S1). Growth rates reported from the literature here are also shown in Figure 5. If a range is given, then the default parameter input to the chemostat model is indicated as a single value beneath the range. See Table S5 for the default parameters for the 3D ocean-biogeochemical model, where the reference values for µ^max^ are 2-fold higher than here to compensate for the modification by temperature.

| Trait | Facultative heterotrophs (NAR, NIR) | Ammonia-oxidizing archaea (AOA) | Nitrite-oxidizing bacteria (NOB) | Anammox bacteria (AOX) |
| --- | --- | --- | --- | --- |
| Model species | Multiple/spectrum (71, 83, 85) | Species belonging to Thaumarchaeota (8, 88) | *Nitrospina* and *Nitrococcus* sp. (8, 10, 13) | *Candidatus Scalindua* sp. (89) |
| Maximum growth rate ($\mu_{\max}$) | 0.01-100 day<sup>-1</sup>; 0.1-1.0 day<sup>-1</sup> on average (71, 90, 91)<br>0.5 day<sup>-1</sup> | ~ 0.2 – 0.9 day<sup>-1</sup> (24, 38, 63–67)<br>0.5 day<sup>-1</sup> | ~ 0.6 – 1.7 day<sup>-1</sup> (24, 68–70)<br>1.0 day<sup>-1</sup> | ~ 0.25 day<sup>-1</sup> (92–94) |
| Cell volume (V) | 0.012 $\mu\text{m}^3$ (90) | †0.025 $\mu\text{m}^3$ (95) | †0.21 $\mu\text{m}^3$ (70) | †0.27 $\mu\text{m}^3$ (96) |
| Biomass stoichiometry | C<sub>4.5</sub>N (97) | C<sub>4</sub>N (80) | C<sub>4</sub>N (80) | C<sub>5</sub>H<sub>8.7</sub>O<sub>1.55</sub>N (92) |
| Cell C quota<sup>a</sup> | 6.5 fg C cell<sup>-1</sup> (97) | 11-17 fg C cell<sup>-1</sup> (80)<br>11.5 fg C cell<sup>-1</sup> | 28-1207 fg C cell<sup>-1</sup> (80)<br>39.8 fg C cell<sup>-1</sup> | 38.5 fg C cell<sup>-1</sup> |
| Biomass yield (y)<sup>β,δ</sup> | 0.05 – 0.6 C<sub>bio</sub> (mol C<sub>org</sub>)<sup>-1</sup> (83)<br>0.31 mol C<sub>bio</sub> (mol C<sub>org</sub>)<sup>-1</sup> (86) | 0.091 ± 0.012 mol C<sub>bio</sub> (mol NH<sub>3</sub>)<sup>-1</sup> (80) | 0.036 ± 0.005 mol C<sub>bio</sub> (mol NO<sub>2</sub>)<sup>-1</sup> (80) | 0.071 ± 0.002 mol C<sub>bio</sub> (mol NH<sub>4</sub>)<sup>-1</sup> (92) |
| Maximum uptake rate ($V_{\max}$)<sup>γ</sup> | 0.96 mol N<sub>org</sub> (mol N<sub>bio</sub>)<sup>-1</sup> day<sup>-1</sup><br>12.7 mol NO<sub>3</sub> (mol N<sub>bio</sub>)<sup>-1</sup> day<sup>-1</sup><br>8.5 mol NO<sub>2</sub> (mol N<sub>bio</sub>)<sup>-1</sup> day<sup>-1</sup> | 20.4 mol NH<sub>4</sub> (mol N<sub>bio</sub>)<sup>-1</sup> day<sup>-1</sup> | 76.2 mol NO<sub>2</sub> (mol N<sub>bio</sub>)<sup>-1</sup> day<sup>-1</sup> | 14.1 mol NH<sub>4</sub> (mol N<sub>bio</sub>)<sup>-1</sup> day<sup>-1</sup> |
| Half-saturation coefficients ($K_s$) | $K_{N_{org}} = 0.1 \mu\text{M}$<br>$K_{NO_x} = 4 \mu\text{M}$ (98, 99) | $K_{NH_4} = 0.065 - 0.133 \mu\text{M}$ NH<sub>4</sub> (63, 65, 100)<br>$K_{NH_4} = 0.1 \mu\text{M}$ | $K_{NO_2} = 0.03 - 0.5 \mu\text{M}$ (101–103)<br>$K_{NO_2} = 0.1 \mu\text{M}$ | $K_{NO_2} = K_{NH_4} = 0.45 \mu\text{M}$ (92, 104) |

<sup>†</sup>Cell volume calculated from dimensions (diameter and length for rods, diameter for cocci).
<sup>α</sup>Unreferenced cell quota for anammox are scaled by cell volume relative to the 6.5 fg C cell<sup>-1</sup> of SAR11 bacteria (97).
<sup>β</sup>Assuming C:N of biomass of 4.5 (97) and a Redfieldian C:N of dissolved organic matter of 7.625 (105).
<sup>δ</sup>Listed yields are those assumed for aerobic heterotrophic metabolisms. Because the relative yields of aerobic vs. anaerobic heterotrophy are unclear, in the model, we assume that the biomass yield of cells carrying out nitrate reduction or denitrification are 10% less than those carrying out aerobic heterotrophy, which represents the underlying energetics, following Zakem et al. (28).
<sup>γ</sup> $V_{\max} = \mu_{\max} / y$ .

**Table S2.**
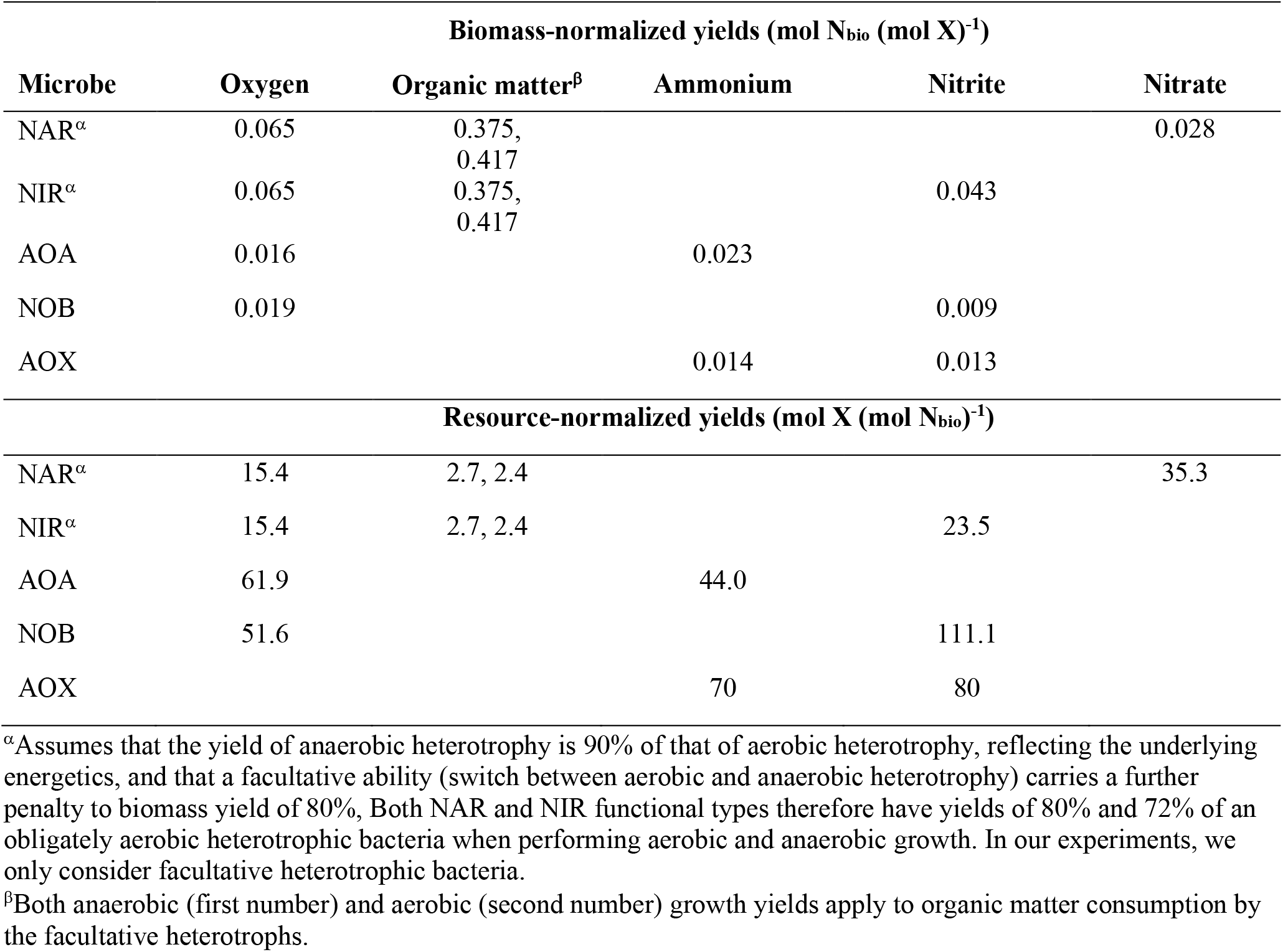
Yields as input to the default chemostat model. These yields are sourced directly from the literature listed in Table S1. Resource-normalized yields are the inverse of the biomass-normalized yields.

| Microbe | Biomass-normalized yields (mol N<sub>bio</sub> (mol X)<sup>-1</sup>) |  |  |  |  |
| --- | --- | --- | --- | --- | --- |
|  | Oxygen | Organic matter<sup>β</sup> | Ammonium | Nitrite | Nitrate |
| NAR<sup>α</sup> | 0.065 | 0.375,<br>0.417 |  |  | 0.028 |
| NIR<sup>α</sup> | 0.065 | 0.375,<br>0.417 |  | 0.043 |  |
| AOA | 0.016 |  | 0.023 |  |  |
| NOB | 0.019 |  |  | 0.009 |  |
| AOX |  |  | 0.014 | 0.013 |  |
| Microbe | Resource-normalized yields (mol X (mol N<sub>bio</sub>)<sup>-1</sup>) |  |  |  |  |
|  | Oxygen | Organic matter<sup>β</sup> | Ammonium | Nitrite | Nitrate |
| NAR<sup>α</sup> | 15.4 | 2.7, 2.4 |  |  | 35.3 |
| NIR<sup>α</sup> | 15.4 | 2.7, 2.4 |  | 23.5 |  |
| AOA | 61.9 |  | 44.0 |  |  |
| NOB | 51.6 |  |  | 111.1 |  |
| AOX |  |  | 70 | 80 |  |

<sup>α</sup>Assumes that the yield of anaerobic heterotrophy is 90% of that of aerobic heterotrophy, reflecting the underlying energetics, and that a facultative ability (switch between aerobic and anaerobic heterotrophy) carries a further penalty to biomass yield of 80%. Both NAR and NIR functional types therefore have yields of 80% and 72% of an obligately aerobic heterotrophic bacteria when performing aerobic and anaerobic growth. In our experiments, we only consider facultative heterotrophic bacteria.
<sup>β</sup>Both anaerobic (first number) and aerobic (second number) growth yields apply to organic matter consumption by the facultative heterotrophs.

**Table S3.** Subsistence concentrations of key resources required to support stable biomasses of microbes. Numbers are based on the traits in Table S1, yields in Table S2 and equations in the methods. Single values are those used for the chemostat and biogeochemical models, while ranges presented in the parentheses are those reflecting reasonable variations in key traits. Red indicates the most competitive microbe for a given resource. Assumes a loss rate of 0.05 day^-1^ common to all microbes. Equations detailed in the methods.

| Microbe | Oxygen (nM) | Ammonium ($\mu\text{M}$) | Nitrite ($\mu\text{M}$) |
| --- | --- | --- | --- |
| NAR<sup>a</sup> | 0.38<br>(0.15 – 2.9) |  |  |
| NIR<sup>a</sup> | 0.38<br>(0.15 – 2.9) |  | 0.65<br>(0.65 – 4.0) |
| AOA | 2.6<br>(2.3 – 3.1) | 0.011<br>(0.007 – 0.015) |  |
| NOB | 9.1<br>(7.9 – 10.7) |  | 0.005<br>(0.002 – 0.026) |
| AOX |  | 0.15<br>(0.15 – 1.0) | 0.15<br>(0.15 – 0.83) |

<sup>a</sup>Assumes that the yield of anaerobic heterotrophy is 90% of that of aerobic heterotrophy, and that facultative ability carries a further penalty of 80% efficiency relative to obligately aerobic heterotrophy (e.g., total penalty of 72 % for biomass synthesis from nitrite reduction by NIR).

**Table S4.** Integrated contribution of NOB and AOA to oxygen consumption within the AMZ environment (oxygen < 1 µM) in the default and sensitivity experiments. Range in parentheses represents minimum and maximum values across 365 daily mean values in final year of simulation.

| Experiment | NOB contribution (%) | AOA contribution (%) |
| --- | --- | --- |
| Default | 42.7<br>(37.8 – 49.3) | 8.9<br>(7.9 – 10.2) |
| $\mu_{\text{Het}}^{\text{max}} = 2.0 \text{ day}^{-1}$ at 30°C | 34.0<br>(29.7 – 39.6) | 10.0<br>(9.0 – 11.3) |
| $\mu_{\text{Het}}^{\text{max}} = 2.0 \text{ day}^{-1}$ at 30°C<br>$\mu_{\text{NOB}}^{\text{max}} = 1.0 \text{ day}^{-1}$ at 30°C | 27.2<br>(24.3 – 31.6) | 11.0<br>(10.1 – 12.5) |
| $\mu_{\text{Het}}^{\text{max}} = 2.0 \text{ day}^{-1}$ at 30°C<br>$\mu_{\text{NOB}}^{\text{max}} = 1.0 \text{ day}^{-1}$ at 30°C<br>$\mu_{\text{AOA}}^{\text{max}} = 2.0 \text{ day}^{-1}$ at 30°C | 27.0<br>(24.0 – 31.2) | 13.3<br>(12.0 – 15.2) |

**Table S5.** Default parameters of the 3D ocean-biogeochemical model related to the microbial functional types. Values tested in sensitivity experiments are in parentheses. Note that the µ^max^ (and all rate parameters) of each functional type is strongly dependent on temperature according to 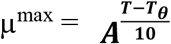. Default values of maximum growth rates 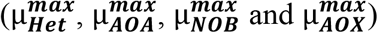 where chosen to be similar to those in the chemostat model (Table S1) at a temperature of 15ºC. Heterotrophs specifically refer to facultative nitrate-reducing and nitrite-reducing types (NAR and NIR; see Fig. 1a).

| Parameter | Description | Value | Units |
| --- | --- | --- | --- |
| C:P | Carbon to phosphorus ratio of heterotrophs and chemoautotrophs | 55:1 | mol mol<sup>-1</sup> |
| C:N | Carbon to nitrogen ratio of heterotrophs and chemoautotrophs | 5:1 | mol mol<sup>-1</sup> |
| Fe:C | Iron to carbon ratio of heterotrophs and chemoautotrophs | 20:1 | μmol mol<sup>-1</sup> |
| $\mu_{Het}^{\max}$ | Maximum growth rate of heterotrophs at $T_{\theta}$ | 1.0 (2.0) | day<sup>-1</sup> |
| $\mu_{AOA}^{\max}$ | Maximum growth rate of AOA at $T_{\theta}$ | 1.0 (2.0) | day<sup>-1</sup> |
| $\mu_{NOB}^{\max}$ | Maximum growth rate of NOB at $T_{\theta}$ | 2.0 (1.0) | day<sup>-1</sup> |
| $\mu_{AOX}^{\max}$ | Maximum growth rate of anammox bacteria at $T_{\theta}$ | 0.5 | day<sup>-1</sup> |
| $A$ | Coefficient for the Arrhenius temperature function | 1.7 | °C<sup>-1</sup> |
| $T_{\theta}$ | Reference temperature | 30 | °C |
| $K_{Het}^{DOC}$ | Half-saturation coefficient for uptake of semi-labile DOC by heterotrophs | 10 | μM |
| $K_{Het}^{DOCr}$ | Half-saturation coefficient for uptake of semi-recalcitrant DOC by heterotrophs | 1000 | μM |
| $K_{Het}^{NO_3}$ | Half-saturation coefficient for uptake of nitrate by heterotrophs | 15 | μM |
| $K_{Het}^{NO_2}$ | Half-saturation coefficient for uptake of nitrite by heterotrophs | 10 | μM |
| $K_{AOA}^{NH_4}$ | Half-saturation coefficient for uptake of ammonium by AOA | 0.1 | μM |
| $K_{NOB}^{NO_2}$ | Half-saturation coefficient for uptake of nitrite by NOB | 0.1 | μM |
| $K_{AOX}^{NH_4}$ | Half-saturation coefficient for uptake of ammonium by anammox bacteria | 0.45 | μM |
| $K_{AOX}^{NO_2}$ | Half-saturation coefficient for uptake of nitrite by anammox bacteria | 0.45 | μM |
| $\rho_{Het}^{O_2}$ | Diffusion-limited oxygen uptake by heterotrophs | 450 | m<sup>3</sup> (μM C<sub>bio</sub>)<sup>-1</sup> day<sup>-1</sup> |
| $\rho_{Het}^{O_2}$ | Diffusion-limited oxygen uptake by AOA | 275 | $\text{m}^3 (\mu\text{M C}_{\text{bio}})^{-1} \text{ day}^{-1}$ |
| $\rho_{Het}^{O_2}$ | Diffusion-limited oxygen uptake by NOB | 67 | $\text{m}^3 (\mu\text{M C}_{\text{bio}})^{-1} \text{ day}^{-1}$ |
| $M^l$ | Linear mortality rate of heterotrophs and chemoautotrophs | $0.05\mu^{max}$ | $\text{day}^{-1}$ |
| $M^q$ | Quadratic mortality rate of heterotrophs and chemoautotrophs | 0.2 | $(\mu\text{M day})^{-1}$ |
| $g_{nZ}^{max}$ | Maximum grazing rate of nano-zooplankton | 4.2 | $\text{day}^{-1}$ |
| $g_Z^{max}$ | Maximum grazing rate of zooplankton | 3.3 | $\text{day}^{-1}$ |
| $K_{nZ}^{p_i}$ | Half-saturation coefficient for grazing of prey ($p_i$) carbon biomass by nano-zooplankton | 6.2 | $\mu\text{M}$ |
| $K_Z^{p_i}$ | Half-saturation coefficient for grazing of prey ($p_i$) carbon biomass by zooplankton | 2.4 | $\mu\text{M}$ |
| $m^{DOM}$ | Fraction of biomass lost via linear and quadratic mortality routed to DOM | 0.5 | |
| $m^{POM}$ | Fraction of biomass lost via linear and quadratic mortality routed to POM | 0.3 | |
| $m^{DIN}$ | Fraction of biomass lost via linear and quadratic mortality routed to dissolved inorganic nutrients ($\text{NH}_4$, DIC, $\text{PO}_4$) | 0.2 | |
| $\varphi^{DOM}$ | Fraction of biomass lost via grazing routed to DOM | 0.06 | |
| $\varphi^{POM}$ | Fraction of biomass lost via grazing routed to POM | 0.0 | |
| $\varphi^{DIN}$ | Fraction of biomass lost via grazing routed to dissolved inorganic nutrients ($\text{NH}_4$, DIC, $\text{PO}_4$) | 0.64 | |
| $\varphi^{DIN}$ | Fraction of biomass lost via grazing routed to grazer biomass | 0.3 | |
| $\varepsilon^{rDOM}$ | Fraction of DOM production routed to semi-recalcitrant pool | 0.3 | |

**Table S6.** Metadata for MAGs. Estimates of µ^max^ were made using a statistical model built from codon usage bias (*50*) at a reference temperature of 13ºC. This statistical model was updated with new growth rate measurements and accompanying genomes of chemoautotrophs (Fig. S13; Table S8). MAGS from the Eastern Tropical South Pacific AMZ are from Sun et al. (*15*), those of NOB from other (oxygenated) regions are collated by Sun et al. (*48*), and those of NOB from the Arabian Sea and Eastern Tropical North Pacific AMZs are from Fortin et al. (*20*).

| Organism | Source | Latitude | Longitude | Depth (m) | # Ribosomal Proteins | Doubling time (hr) | $\mu^{\max}$ |
| --- | --- | --- | --- | --- | --- | --- | --- |
| Pelagibacter spp. | South East Tropical Pacific AMZ | -20.50 | -70.70 | 45, 80, 200 and 300 | 23 | 27.0 | 0.61 |
| Pelagibacter spp. | South East Tropical Pacific AMZ | -20.50 | -70.70 | 45, 80, 200 and 300 | 12 | 35.6 | 0.47 |
| Pelagibacter spp. | South East Tropical Pacific AMZ | -20.50 | -70.70 | 45, 80, 200 and 300 | 20 | 20.6 | 0.81 |
| Unknown sp. in Nitrosopumilus genus | South East Tropical Pacific AMZ | -20.50 | -70.70 | 45, 80, 200 and 300 | 27 | 14.1 | 1.18 |
| Nitrosopumilus sp. 002730325 | South East Tropical Pacific AMZ | -20.50 | -70.70 | 45, 80, 200 and 300 | 27 | 12.7 | 1.31 |
| Unknown sp. in Nitrosopelagicus genus | South East Tropical Pacific AMZ | -20.50 | -70.70 | 45, 80, 200 and 300 | 20 | 15.1 | 1.10 |
| Nitrosopumilus sp002690535 | South East Tropical Pacific AMZ | -20.50 | -70.70 | 45, 80, 200 and 300 | 32 | 18.6 | 0.89 |
| ETSP MAG-1 in Nitrospinaceae family | South East Tropical Pacific AMZ | -20.50 | -70.70 | 45, 80, 200 and 300 | 37 | 9.1 | 1.84 |
| ETSP MAG-2 in Nitrospinaceae family | South East Tropical Pacific AMZ | -20.50 | -70.70 | 45, 80, 200 and 300 | 41 | 9.6 | 1.73 |
| Nitrospina sp. (AB-629-B18) | North Atlantic Gyre | 18.17 | -41.03 | 511 | 47 | 10.3 | 1.62 |
| Nitrospina sp. (AG-538-K21) | North Atlantic Gyre | 18.17 | -41.03 | 511 | 30 | 8.6 | 1.92 |
| Nitrospina sp. (AG-538-L21) | North Atlantic Gyre | 18.17 | -41.03 | 511 | 35 | 7.6 | 2.19 |
| Nitrospina sp. (Bin 24 Ga0113616) | Northern Gulf of Mexico | 28.17 | 90.83 | 16 | 43 | 13.5 | 1.23 |
| Nitrospina sp. (Bin 25 Ga0113617) | Northern Gulf of Mexico | 28.17 | 90.83 | 16 | 43 | 11.1 | 1.50 |
| Nitrospina sp. (Bin 47 2 Ga0113620) | Northern Gulf of Mexico | 28.17 | 90.83 | 16 | 39 | 11.1 | 1.50 |
| Nitrospina sp. LS-NOB | Sponge metagenome from Lophophysema sp. RT-2015 collected from seamount in<br>South China Sea | N/A | N/A | N/A | 40 | 12.8 | 1.23 |
| Nitrospina sp.<br>NAT221 | Marine pelagic<br>biome (5-1000 m) | N/A | N/A | N/A | 35 | 8.1 | 2.07 |
| Nitrospina<br>gracilis 3/211 | Culture | N/A | N/A | N/A | 41 | 13.8 | 1.21 |
| Nitrospina sp.<br>UBA6737 | Deep-sea sponge<br>Neamphius huxleyi | N/A | N/A | N/A | 39 | 12.9 | 1.29 |
| NOB1a in<br>Nitrospinaceae<br>family | Eastern Tropical<br>North Pacific AMZ | 18.0 | -102.0 | 70 | 40 | 10.0 | 1.67 |
| NOB1b in<br>Nitrospinaceae<br>family | Eastern Tropical<br>North Pacific AMZ | 18.0 | -102.0 | 70 | 40 | 10.3 | 1.62 |
| NOB1c in<br>Nitrospinaceae<br>family | Eastern Tropical<br>North Pacific AMZ | 16.0 | -105.0 | 140 | 38 | 10.1 | 1.65 |
| NOB1d in<br>Nitrospinaceae<br>family | Eastern Tropical<br>North Pacific AMZ | 16.0 | -105.0 | 130 | 39 | 10.2 | 1.63 |
| NOB1e in<br>Nitrospinaceae<br>family | Eastern Tropical<br>North Pacific AMZ | 18.0 | -102.0 | 45 | 40 | 9.7 | 1.72 |
| NOB1f in<br>Nitrospinaceae<br>family | Arabian Sea AMZ | 15.15 | 64.0 | 130 | 39 | 9.2 | 1.81 |
| NOB1g in<br>Nitrospinaceae<br>family | Arabian Sea AMZ | 15.15 | 64.0 | 150 | 39 | 9.0 | 1.85 |
| NOB1h in<br>Nitrospinaceae<br>family | Arabian Sea AMZ | 15.15 | 64.0 | 200 | 40 | 9.2 | 1.81 |
| NOB1i in<br>Nitrospinaceae<br>family | Arabian Sea AMZ | 15.15 | 64.0 | 400 | 46 | 8.9 | 1.87 |
| NOB2a in<br>Nitrospinaceae<br>family | Eastern Tropical<br>North Pacific AMZ | 18.7 | -104.4 | 200 | 40 | 10.1 | 1.65 |
| NOB2b in<br>Nitrospinaceae<br>family | Eastern Tropical<br>North Pacific AMZ | 18.7 | -104.4 | 750 | 37 | 9.8 | 1.69 |
| NOB2c in<br>Nitrospinaceae<br>family | Eastern Tropical<br>North Pacific AMZ | 18.0 | -102.0 | 800 | 40 | 10.2 | 1.63 |
| NOB2d in<br>Nitrospinaceae<br>family | Eastern Tropical<br>North Pacific AMZ | 17.2 | -110.7 | 185 | 39 | 10.6 | 1.57 |
| NOB2e in<br>Nitrospinaceae<br>family | Eastern Tropical<br>North Pacific AMZ | 17.2 | -110.7 | 215 | 39 | 11.3 | 1.47 |
| NOB3a in<br>Nitrospinaceae<br>family | Eastern Tropical<br>North Pacific AMZ | 18.7 | -104.4 | 750 | 37 | 12.1 | 1.38 |
| NOB3b in<br>Nitrospinaceae<br>family | Eastern Tropical<br>North Pacific AMZ | 18.0 | -102.0 | 800 | 39 | 11.5 | 1.44 |
| NOB3c in<br>Nitrospinaceae<br>family | Eastern Tropical<br>North Pacific AMZ | 16.0 | -105.0 | 850 | 39 | 12.1 | 1.37 |
| NOB3d in Nitrospinaceae family | Arabian Sea AMZ | 15.15 | 64.0 | 150 | 34 | 10.7 | 1.56 |
| NOB4a in Nitrospinaceae family | Eastern Tropical North Pacific AMZ | 17.2 | -110.7 | 185 | 43 | 14.2 | 1.17 |
| NOB4b in Nitrospinaceae family | Eastern Tropical North Pacific AMZ | 16.0 | -105.0 | 140 | 43 | 14.0 | 1.19 |
| NOB5a in Nitrospinaceae family | Eastern Tropical North Pacific AMZ | 16.0 | -105.0 | 130 | 41 | 14.6 | 1.14 |
| NOB5b in Nitrospinaceae family | Eastern Tropical North Pacific AMZ | 16.0 | -105.0 | 140 | 41 | 15.3 | 1.09 |
| NOB6a in Nitrospinaceae family | Eastern Tropical North Pacific AMZ | 16.0 | -105.0 | 130 | 19 | 12.5 | 1.33 |
| NOB6b in Nitrospinaceae family | Arabian Sea AMZ | 15.15 | 64.0 | 130 | 25 | 14.4 | 1.16 |
| NOB7a in Nitrospinaceae family | Arabian Sea AMZ | 15.15 | 64.0 | 150 | 41 | 14.1 | 1.18 |
| NOB8a in Nitrospinaceae family | Arabian Sea AMZ | 15.15 | 64.0 | 150 | 42 | 14.7 | 1.13 |
| NOB8b in Nitrospinaceae family | Arabian Sea AMZ | 15.15 | 64.0 | 400 | 18 | 14.0 | 1.19 |
| NOB9a in Nitrospinaceae family | Arabian Sea AMZ | 15.15 | 64.0 | 200 | 21 | 18.9 | 0.88 |
| NOB9b in Nitrospinaceae family | Arabian Sea AMZ | 15.15 | 64.0 | 400 | 33 | 16.9 | 0.98 |
| NOB10a in Nitrospinaceae family | Arabian Sea AMZ | 15.15 | 64.0 | 400 | 38 | 14.5 | 1.15 |
| NOB10b in Nitrospinaceae family | Arabian Sea AMZ | 15.15 | 64.0 | 200 | 39 | 13.4 | 1.24 |

**Table S7.** Metadata for AOA and NOB used to train the new version of gRodon. Performance of the new training of gRodon is presented in Fig. S13.

| Species | Doubling time (hours) | OGT (°C) | Notes | Reference | Genome | Type |
| --- | --- | --- | --- | --- | --- | --- |
| *Nitrososphaera viennensis* | 29.2 | 45 | Isolate, soil | (106) | GCF_000698785.1 | AOA |
| *Candidatus Nitrosocosmicus hydrocola* | 51.7 | 33 | Enrichment culture (99%), wastewater treatment plant | (107) | GCF_001870125.1 | AOA |
| *Nitrosopumilus maritimus SCM1* | 21.32 | 28 | Isolate, marine | (67) | GCF_000018465.1 | AOA |
| *Nitrolancea hollandica* | 36.16 | 40 | Isolate from bioreactor | (108) | GCF_000297255.1 | NOB |
| *Candidatus Nitrotoga arctica* | 54 | 16 | Soil, enrichment | (109) | GCF_918378365.1 | NOB |
| *Nitrococcus mobilis* | 9.79 | 28 | Marine | (110) | GCF_000153205.1 | NOB |
| *Nitrospina gracilis* | 23.77 | 28 | Marine | (52) | GCF_000341545.2 | NOB |
| *Nitrospinae watsonii* | 23.1 | 28 | Isolate, marine | (70) | GCF_946900835.1 | NOB |
| *Nitrospira moscoviensis* | 12 | 39 | Isolate, heating pipe | (111) | GCF_001273775.1 | NOB |

**Table S8.**
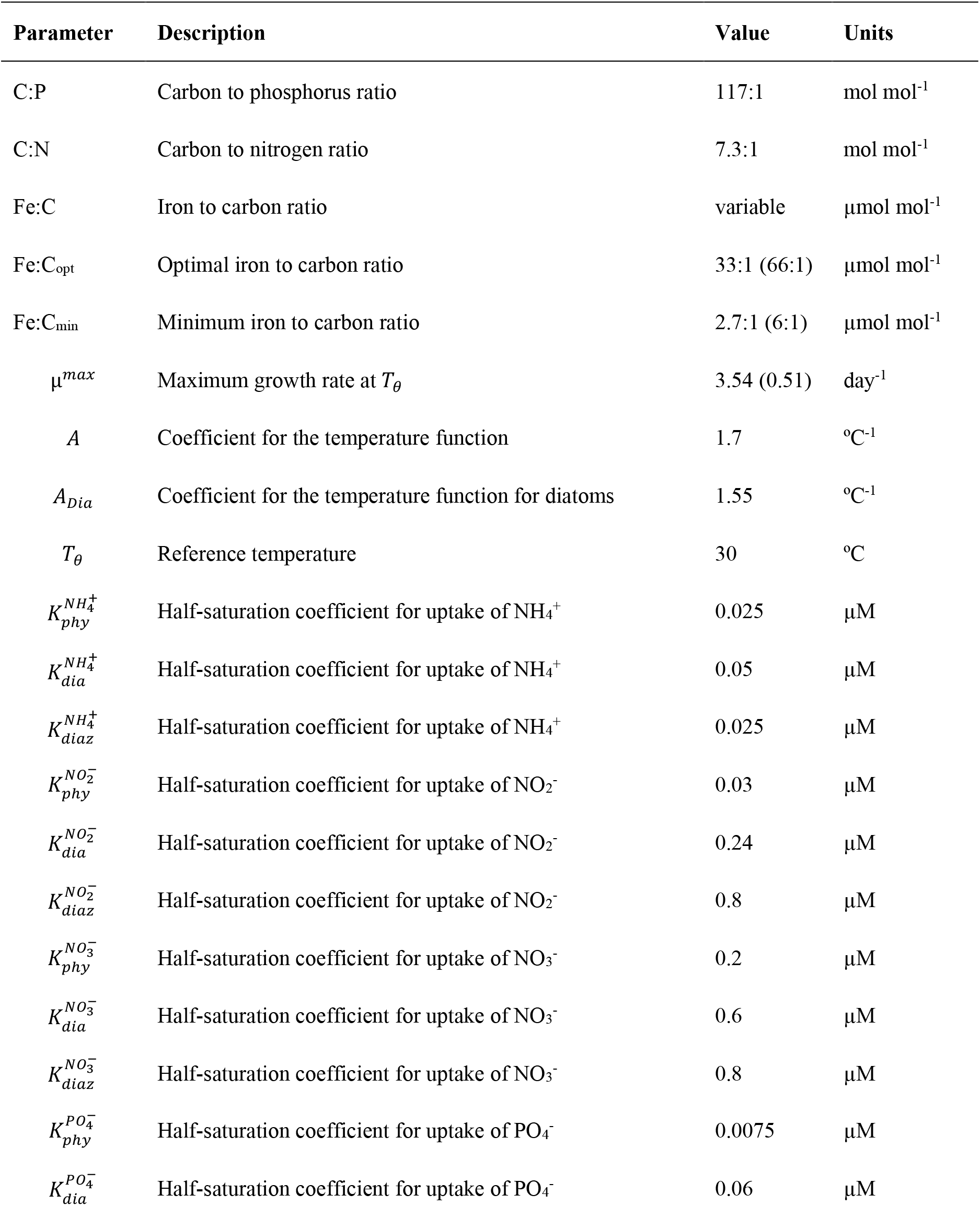
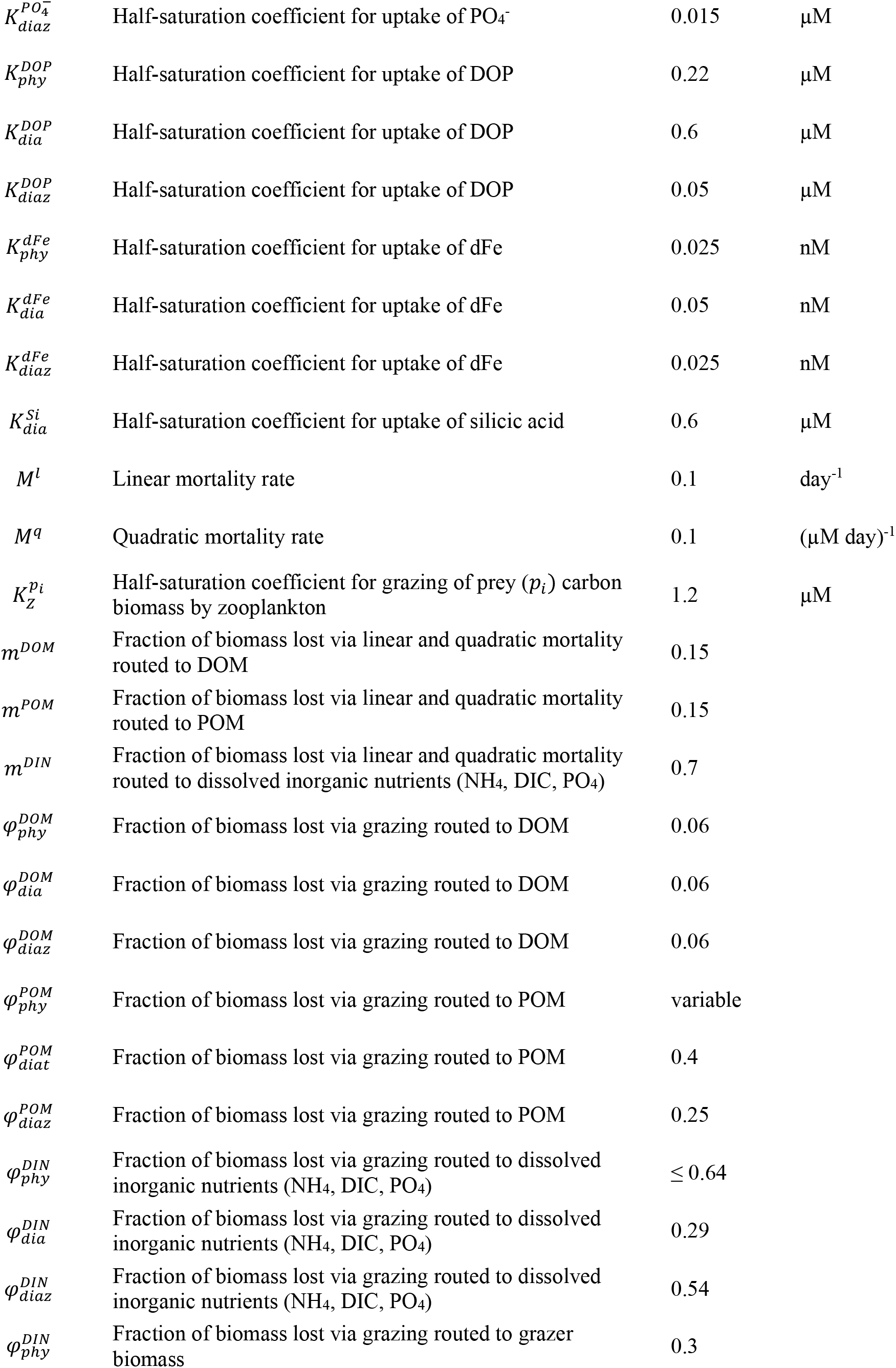
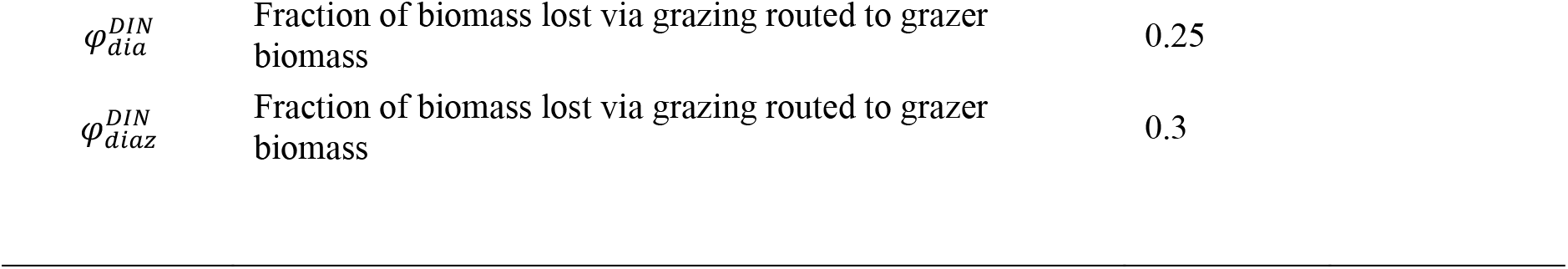
Parameters of the ocean-biogeochemical model related to the phytoplankton functional types. Phy = nanophytoplankton. Dia = diatoms. Diaz = facultative diazotrophs. Diazotrophic phytoplankton type in parentheses.

| Parameter | Description | Value | Units |
| --- | --- | --- | --- |
| C:P | Carbon to phosphorus ratio | 117:1 | mol mol<sup>-1</sup> |
| C:N | Carbon to nitrogen ratio | 7.3:1 | mol mol<sup>-1</sup> |
| Fe:C | Iron to carbon ratio | variable | μmol mol<sup>-1</sup> |
| Fe:C<sub>opt</sub> | Optimal iron to carbon ratio | 33:1 (66:1) | μmol mol<sup>-1</sup> |
| Fe:C<sub>min</sub> | Minimum iron to carbon ratio | 2.7:1 (6:1) | μmol mol<sup>-1</sup> |
| $\mu^{max}$ | Maximum growth rate at $T_\theta$ | 3.54 (0.51) | day<sup>-1</sup> |
| $A$ | Coefficient for the temperature function | 1.7 | °C<sup>-1</sup> |
| $A_{Dia}$ | Coefficient for the temperature function for diatoms | 1.55 | °C<sup>-1</sup> |
| $T_\theta$ | Reference temperature | 30 | °C |
| $K_{phy}^{NH_4^+}$ | Half-saturation coefficient for uptake of NH<sub>4</sub><sup>+</sup> | 0.025 | μM |
| $K_{dia}^{NH_4^+}$ | Half-saturation coefficient for uptake of NH<sub>4</sub><sup>+</sup> | 0.05 | μM |
| $K_{diaz}^{NH_4^+}$ | Half-saturation coefficient for uptake of NH<sub>4</sub><sup>+</sup> | 0.025 | μM |
| $K_{phy}^{NO_2^-}$ | Half-saturation coefficient for uptake of NO<sub>2</sub><sup>-</sup> | 0.03 | μM |
| $K_{dia}^{NO_2^-}$ | Half-saturation coefficient for uptake of NO<sub>2</sub><sup>-</sup> | 0.24 | μM |
| $K_{diaz}^{NO_2^-}$ | Half-saturation coefficient for uptake of NO<sub>2</sub><sup>-</sup> | 0.8 | μM |
| $K_{phy}^{NO_3^-}$ | Half-saturation coefficient for uptake of NO<sub>3</sub><sup>-</sup> | 0.2 | μM |
| $K_{dia}^{NO_3^-}$ | Half-saturation coefficient for uptake of NO<sub>3</sub><sup>-</sup> | 0.6 | μM |
| $K_{diaz}^{NO_3^-}$ | Half-saturation coefficient for uptake of NO<sub>3</sub><sup>-</sup> | 0.8 | μM |
| $K_{phy}^{PO_4^-}$ | Half-saturation coefficient for uptake of PO<sub>4</sub><sup>-</sup> | 0.0075 | μM |
| $K_{dia}^{PO_4^-}$ | Half-saturation coefficient for uptake of PO<sub>4</sub><sup>-</sup> | 0.06 | μM |
| $K_{diaz}^{PO_4^-}$ | Half-saturation coefficient for uptake of $PO_4^-$ | 0.015 | $\mu M$ |
| $K_{phy}^{DOP}$ | Half-saturation coefficient for uptake of DOP | 0.22 | $\mu M$ |
| $K_{dia}^{DOP}$ | Half-saturation coefficient for uptake of DOP | 0.6 | $\mu M$ |
| $K_{diaz}^{DOP}$ | Half-saturation coefficient for uptake of DOP | 0.05 | $\mu M$ |
| $K_{phy}^{dFe}$ | Half-saturation coefficient for uptake of dFe | 0.025 | nM |
| $K_{dia}^{dFe}$ | Half-saturation coefficient for uptake of dFe | 0.05 | nM |
| $K_{diaz}^{dFe}$ | Half-saturation coefficient for uptake of dFe | 0.025 | nM |
| $K_{dia}^{Si}$ | Half-saturation coefficient for uptake of silicic acid | 0.6 | $\mu M$ |
| $M^l$ | Linear mortality rate | 0.1 | $day^{-1}$ |
| $M^q$ | Quadratic mortality rate | 0.1 | $(\mu M \text{ day})^{-1}$ |
| $K_z^{p_i}$ | Half-saturation coefficient for grazing of prey ($p_i$) carbon biomass by zooplankton | 1.2 | $\mu M$ |
| $m^{DOM}$ | Fraction of biomass lost via linear and quadratic mortality routed to DOM | 0.15 | |
| $m^{POM}$ | Fraction of biomass lost via linear and quadratic mortality routed to POM | 0.15 | |
| $m^{DIN}$ | Fraction of biomass lost via linear and quadratic mortality routed to dissolved inorganic nutrients ($NH_4$, DIC, $PO_4$) | 0.7 | |
| $\phi_{phy}^{DOM}$ | Fraction of biomass lost via grazing routed to DOM | 0.06 | |
| $\phi_{dia}^{DOM}$ | Fraction of biomass lost via grazing routed to DOM | 0.06 | |
| $\phi_{diaz}^{DOM}$ | Fraction of biomass lost via grazing routed to DOM | 0.06 | |
| $\phi_{phy}^{POM}$ | Fraction of biomass lost via grazing routed to POM | variable | |
| $\phi_{diat}^{POM}$ | Fraction of biomass lost via grazing routed to POM | 0.4 | |
| $\phi_{diaz}^{POM}$ | Fraction of biomass lost via grazing routed to POM | 0.25 | |
| $\phi_{phy}^{DIN}$ | Fraction of biomass lost via grazing routed to dissolved inorganic nutrients ($NH_4$, DIC, $PO_4$) | $\leq 0.64$ | |
| $\phi_{dia}^{DIN}$ | Fraction of biomass lost via grazing routed to dissolved inorganic nutrients ($NH_4$, DIC, $PO_4$) | 0.29 | |
| $\phi_{diaz}^{DIN}$ | Fraction of biomass lost via grazing routed to dissolved inorganic nutrients ($NH_4$, DIC, $PO_4$) | 0.54 | |
| $\phi_{phy}^{DIN}$ | Fraction of biomass lost via grazing routed to grazer biomass | 0.3 | |

|  |  |  |
| --- | --- | --- |
| $\varphi_{dia}^{DIN}$ | Fraction of biomass lost via grazing routed to grazer biomass | 0.25 |
| $\varphi_{diaz}^{DIN}$ | Fraction of biomass lost via grazing routed to grazer biomass | 0.3 |

**Movie S1**.

Animation of oxygen concentrations in the 3D ROMS-BEC2 model with explicit microbial functional types at daily resolution over the final year of the default 51-year simulation. Left panel is average between 200-400 meters. Right panel is a latitudinal section along 85ºW.

**Movie S2**.

Animation of nitrite concentrations in the 3D ROMS-BEC2 model with explicit microbial functional types at daily resolution over the final year of the default 51-year simulation. Left panel is average between 200-400 meters. Right panel is a latitudinal section along 85ºW.

**Movie S3**.

Animation of ammonia oxidation rate in the 3D ROMS-BEC2 model with explicit microbial functional types at daily resolution over the final year of the default 51-year simulation. Left panel is the maximum between 200-400 meters. Right panel is a latitudinal section along 85ºW.

**Movie S4**.

Animation of nitrite oxidation rate in the 3D ROMS-BEC2 model with explicit microbial functional types at daily resolution over the final year of the default 51-year simulation. Left panel is the maximum between 200-400 meters. Right panel is a latitudinal section along 85ºW.

**Movie S5**.

Animation of heterotrophic nitrate reduction in the 3D ROMS-BEC2 model with explicit microbial functional types at daily resolution over the final year of the default 51-year simulation. Left panel is the maximum between 200-400 meters. Right panel is a latitudinal section along 85ºW.

**Movie S6**.

Animation of heterotrophic nitrite reduction in the 3D ROMS-BEC2 model with explicit microbial functional types at daily resolution over the final year of the default 51-year simulation. Left panel is the maximum between 200-400 meters. Right panel is a latitudinal section along 85ºW.

**Movie S7**.

Animation of anammox in the 3D ROMS-BEC2 model with explicit microbial functional types at daily resolution over the final year of the default 51-year simulation. Left panel is the maximum between 200-400 meters. Right panel is a latitudinal section along 85ºW.

**Movie S8**.

Animation of the ratio of NOB to AOA in the 3D ROMS-BEC2 model with explicit microbial functional types at daily resolution over the final year of the default 51-year simulation. Left panel is average between 200-400 meters. Right panel is a latitudinal section along 85ºW.

**Movie S9**.

Animation of the contribution of NOB to oxygen consumption in the 3D ROMS-BEC2 model with explicit microbial functional types at daily resolution over the final year of the default 51-year simulation. Left panel is the maximum between 200-400 meters. Right panel is a latitudinal section along 85ºW.

## Notes

https://doi.org/10.5281/zenodo.15137773

https://doi.org/10.5281/zenodo.15139207

https://doi.org/10.5281/zenodo.15137671

https://doi.org/10.5281/zenodo.15139307

### Competing Interest Statement

The authors have declared no competing interest.

### Summary of Updates

- edited the manuscript and figures for improved clarity regarding the novelty of our conclusions - expanded the sensitivity experiments, now using the 3D model, to explore the contribution of nitrite oxidation to oxygen consumption considering uncertainties in the maximum growth rates of the functional types. Using these experiments, we now report a range in the contribution (24-49%; Fig. 4). These tests also demonstrate the robustness of our main conclusion. - included a more thorough analysis of nitrification rates and abundances of NOB and AOA both within and without anoxic zones (Fig. 3). - expanded the analysis of the metagenome-based ("gRodon") estimates of the maximum growth rates. We added data from cultures to the training dataset and compare the resulting estimates to measurements (Fig. 5). - moved the gRodon section to later in the paper to clarify that these estimates serve as a test of our model hypothesis, rather than as input to the models.

